# Luminance invariant encoding in primary visual cortex

**DOI:** 10.1101/2024.04.18.590073

**Authors:** Ronan O’Shea, Ian Nauhaus, Xue-Xin Wei, Nicholas J. Priebe

## Abstract

The retina maintains sensitivity over a large range of luminance intensities by switching between rod and cone photoreceptors. This luminance adaptation has been shown to alter the receptive fields and interneuronal correlations of retinal ganglion cells (RGCs). While these adaptations allow the retina to encode visual information across environmental conditions, they present a challenge to downstream processing areas for which it is important that representations are invariant to light level. We measured the effects of scotopic versus photopic luminance adaptation on thalamic and cortical activity by tracking neuronal populations across light levels. While changes in the output of the retina are evident in the lateral geniculate nucleus (LGN), the representation in primary visual cortex (V1) is largely invariant to the changes in luminance. We show that an invariant V1 code can emerge through the integration of parallel functional pathways at the geniculocortical synapse.

## Introduction

Each day, we traverse disparate sensory environments-from hot midday sun to cool starlit night, loud barrooms and silent hallways, all the while maintaining sensitivity to the world around us. This feat is accomplished by adaptation of neural responses to changes in environmental statistics. This adaptation enables the mammalian visual system to operate across a massive range of environmental drive, with light levels varying over 10 orders of magnitude between midnight and midday. A key component of luminance adaptation is the shift in sensory receptors themselves: from rod to cone photoreceptors (Shapley and Enroth-Cugell, 1984). Despite the shift in which sensory receptors are used to transduce light, mammals can perform a wide array of visually guided tasks in both the scotopic (dim) and photopic (bright) regimes (Maier and White, 1998; Kotler et al., 2010; Rahlfs and Fichtel, 2010; Falcy and Danielson, 2013; Bennie et al., 2014; Dickman and Newsome, 2015; Robbers et al., 2015; Yiu et al., 2022). Here we examine how the neural code in the visual system is affected by, and remains robust to, changes at the periphery.

One strategy that the visual system could take is to generate a representation of the visual world that is invariant to absolute luminance. A major challenge to this strategy is the shifting response properties of the retinal output between rod and cone-mediated vision. In the canonical view of scotopic vs photopic vision, the signal relayed by the retinal ganglion cells (RGCs) varies significantly with luminance. Temporal integration is longer in scotopic conditions (Enroth-Cugell and Shapley, 1973) and spatial receptive field centers grow as surrounds weaken at scotopic luminance levels (Barlow et al., 1957). These changes have the benefit of increasing spatiotemporal integration of photons in a low signal to noise ratio regime, but also present a serious challenge to invariant encoding of visual scenes by downstream regions as the spatiotemporal features encoded by the spiking of RGCs varies with luminance.

A potential mechanism for mitigating the downstream impact of these light dependent changes in retinal coding is proportional adaptation of RGC responses across cell classes (Ruda et al., 2022). That is, while the receptive field properties of RGCs of a given type change predictably with luminance, the relation between these receptive field properties across types remains constant. If downstream neurons integrate functionally diverse retinal inputs, representations of the visual scene could rely less on light level. Indeed, despite changes in RGC spatial tuning, neurons in macaque V1 exhibit invariant orientation and size tuning across light levels (Duffy and Hubel, 2007). In addition, spatial frequency tuning in mouse V1 is well matched under rod versus cone dominated regimes (Rhim et al., 2021).

In addition to changes in the receptive field structure of RGCs, the relationship between neuronal responses also shifts with light level: in scotopic conditions, neighboring RGCs of the same type generally exhibit larger correlated variability, or “noise correlations”, than in photopic conditions (Greschner et al., 2011; Ruda et al., 2020). Changes in the noise correlation structure of RGCs could limit information transmission, particularly if correlated noise in neural responses varies along the same dimensions as the signal (Moreno-Bote et al., 2014). A population decoder of RGC activity which assumes independent variability across neurons fails on scotopic data, but performs well when the full covariance matrix of the population is employed (Ruda et al., 2020). This result suggests that the larger noise correlations between RGCs under scotopic conditions could significantly alter encoding relative to the lower noise correlation photopic state. It is possible that such changes in the population noise may persist in downstream neural populations and limit information transmission at scotopic luminance levels.

To determine how the neural code in central visual areas shifts across light levels, we measured population activity in the visual thalamus and cortex. Using a combination of two-photon calcium imaging and electrophysiology techniques, we find that the evoked responses of the V1 population to diverse stimuli are remarkably consistent between scotopic and photopic conditions. In contrast, we find that their thalamic inputs exhibit substantial shifts in both spatial representations and noise correlations with changes in luminance. A simple model based on the integration of functionally diverse thalamic inputs demonstrates how a convergent architecture acts to maintain a constant representation of the visual world despite the changing structure of afferent input across light levels.

## Results

### Trial-averaged responses in V1 are highly correlated in scotopic and photopic conditions

To examine how population activity in V1 changes with light level we recorded the population activity of CaMKII+ excitatory neurons using two-photon fluorescent calcium imaging in awake mouse V1. Animals viewed natural movies or static gratings at scotopic and photopic luminance levels (Fig. 1a), presented in the lower visual field (anterior V1). The anterior portion of V1 receives retinal input from M-opsin expressing cones under photopic conditions which have a similar chromatic response profile to rods in the mouse retina (Wang et al., 2011; Chang et al., 2013; Rhim et al., 2017,2021). The similarity in chromatic response profiles allows us to use the same chromatic stimulus (525nm) for both scotopic and photopic light levels, varying only the luminance of the stimulus (Fig. 1a).

**Figure 1.**
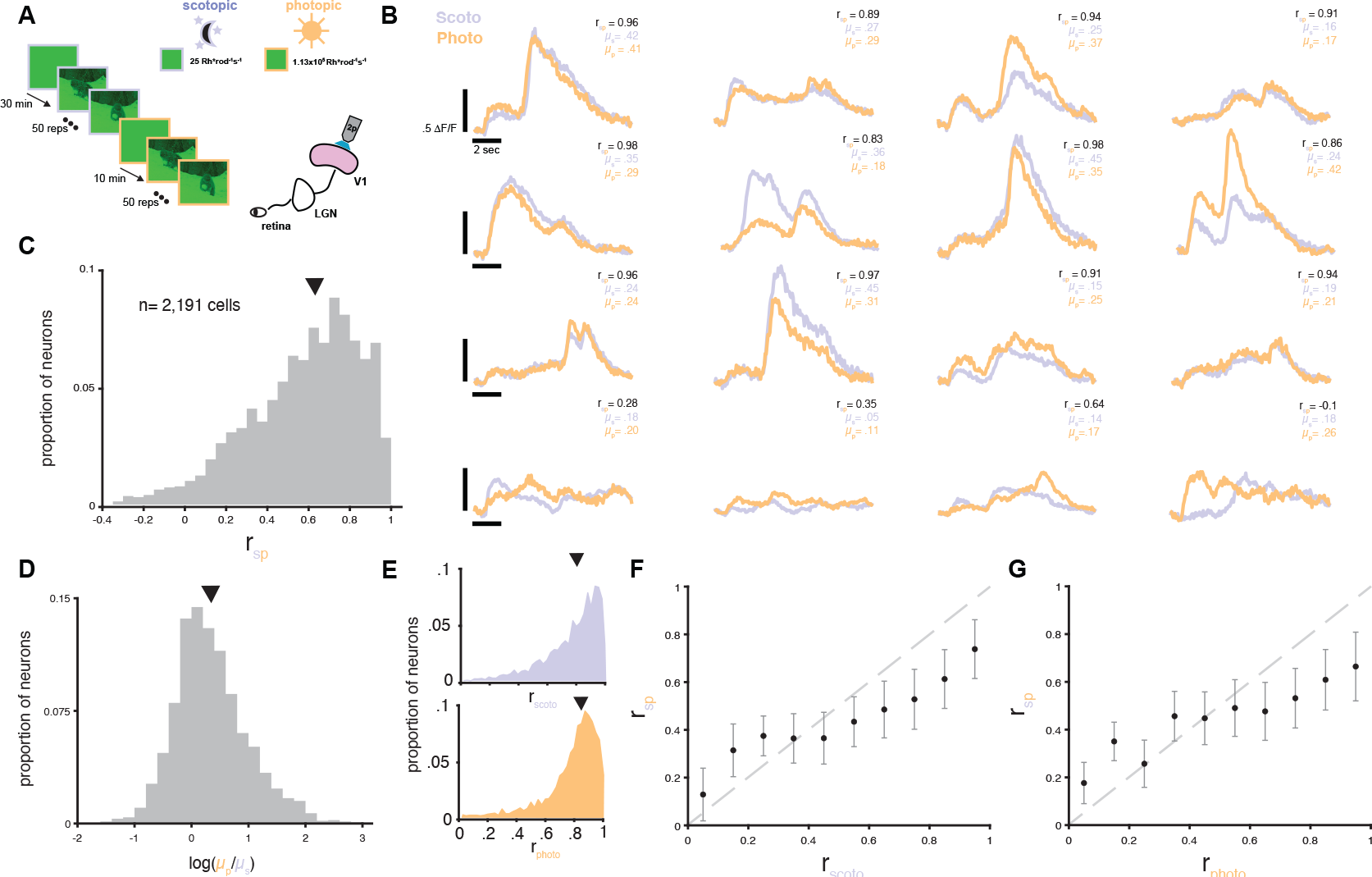
V1 responses to natural scenes in scotopic and photopic conditions. A. Experimental design. Awake mice were head fixed on a treadmill underneath a two-photon microscope while viewing 50 repeats of a 10-second natural movies presented under scotopic (purple) and photopic (orange) conditions. To compare V1 responses in these two regimes, the same V1 neurons were tracked across conditions. B. Trial-averaged, baseline normalized fluorescent responses (F/F) of 16 example cells to a movie at scotopic (purple) and photopic (orange) light levels. Inset gives the cross correlation of the mean responses across light conditions (r_sp_) and the mean over time of the baseline-normalized responses in scotopic (╭_s_) and photopic (╭_p_) conditions. The top 3 rows show example cell responses with high r_sp_. The bottom row shows example cell responses with low r_sp_. C. Histogram of r_sp_ for all cells tracked across luminance conditions. Arrow indicates median of distribution. D. Histogram of the log ratio of ╭_p_ /╭_s_ for all cells tracked across luminance conditions. Arrow indicates median of distribution. E. Histograms for within-condition correlation for trial-averaged responses taken from random splits of trials in scotopic (r_scoto_) (top) and photopic (r_photo_) (bottom) conditions. Arrows indicate median of distributions. F. r_sp_ plotted as function of r_scoto_, binned by within-condition correlation in 10 equally sized bins spanning 0 to 1. Scatter points indicate mean r_sp_ within each bin, bars indicate +/-.5 standard deviation. Gray dashed line indicates unity. G. r_sp_ plotted as function of r_photo_, binned by within-condition correlation in 10 equally sized bins spanning 0 to 1. Scatter points indicate mean r_sp_ within each bin, bars indicate +/-.5 standard deviation. Gray dashed line indicates unity.

We first measured how the mean evoked activity of individual V1 neurons changed across scotopic and photopic conditions in response to natural movies. Despite the known changes in the spatiotemporal response properties of RGCs across luminance levels, we observed a remarkably high correlation in the trial-averaged time courses of responses between the two conditions (Fig. 1b-c). For 2191 visually responsive cells across 4 animals in 10 sessions (mean=219 +/-153 cells/session), the median scotopic-photopic correlation in the trial-averaged time course for single cells (r_sp_) was .62 (Fig. 1c). Responses to the photopic stimulus (╭_p_) were significantly larger than scotopic responses (╭_s_) (photopic median = .147 ΔF/F, scotopic median=.107 ΔF/F, p<.001 two-sided Wilcoxon rank sum test). 69% of cells showed an increase in response amplitude in the photopic relative to scotopic condition, with a median change of +30% (Fig 1d). This is difference in response amplitude is consistent with records in rodent RGCs demonstrating higher firing rates under photopic relative to scotopic conditions (Ruda et al., 2020).

The correlation between responses at these two light levels could be limited both by receptive field changes in V1 due to changes in the retinal input but also by response variability that is independent of light adaptation state (Sahani and Linden, 2003). To quantify response variability within each luminance condition, we repeatedly split trials randomly within each condition and computed the correlation in mean responses for the two pseudo-groups. Correlation computed in this way gives a measure of the reliability of each cell’s response to an identical stimulus at constant luminance. The median within-condition correlation was .82 for scotopic trials (r_scoto_) and .83 for photopic trials (r_photo_) (Fig 1e), indicating that for most neurons, responses were reliable. Importantly, there was a strong correlation between r_scoto_ and r_photo_ for single cells (Pearson’s correlation coefficient = .31), indicating that that trial-to-trial variability in single cell responses remains constant across light conditions. There was a strong, positive relationship between the within-condition and cross condition (r_sp_) correlation for each cell (Fig 1f, g), suggesting that r_sp_ is limited, to some extent, by trial-to-trial variability that is independent of luminance.

If the slope of the relationship between the within-condition correlation and r_sp_ were equal to 1, we would conclude that light condition does not affect the average evoked response in V1. However, a slope of less than 1, as we see in Fig. 1f, g, indicates that changes in the rod versus cone–mediated input do affect the V1 response. To quantify the effect of changing retinal input, we normalized r_sp_ by the within-condition correlation. If the measured cross-condition correlation were limited only by luminance-independent noise this normalized metric should approach 1. Normalizing r_sp_ by r_scoto_ results in a median correlation of .75; normalizing by r_photo_ results in a median correlation of .72, indicating that responses are largely consistent across light level.

To test whether the consistency of V1 responses across light levels generalize to other stimuli, we performed the same analyses using static gratings instead of natural movies. Static oriented gratings were presented at 2 contrasts (30%, 100%), under scotopic and photopic conditions (Supplementary Fig. 1a). V1 single-cell responses to static gratings were highly consistent across luminance conditions, with a median r_sp_ of .77 (n = 14,320, Supplementary Fig. 1 b, c). Trial-averaged response magnitude was higher for the photopic than scotopic conditions (photopic median = .049 ΔF/F, scotopic median=.046 ΔF/F) (Supplementary Fig. 1d). The consistency of within-condition responses to the gratings was highly predictive of r_sp_ (Supplementary Fig. 1f, g). Normalizing r_sp_ by r_scoto_ results in a median correlation of .88; normalizing by r_photo_ results in a median correlation of .86. Therefore, the degree of consistency which we find in V1 responses across light levels is common to natural movie and static grating stimuli.

Our calcium imaging records exhibit high correlations in the evoked responses of V1 cells to natural movies and gratings across light levels, which suggests that the spatiotemporal tuning properties of these cells are also consistent across light levels. We directly measured these tuning properties by presenting drifting gratings over a range of spatial and temporal frequencies at scotopic and photopic light levels while densely sampling V1 single unit activity with the Neuropixels probes (Jun et al., 2017). We tracked single units across light levels and used their trial-averaged firing rates for each stimulus to fit Gaussians to their spatial and temporal frequency responses (Methods, Fig. 2a, d). Using the Center-of-Mass (CoM) of these tuning curves to compare selectivity, we did not find a shift of the distribution of spatial frequency tuning in the V1 population across the two light levels (scotopic CoM mean= .042 +/-.022 cyc/deg, photopic CoM mean= .042 +/-.022 cyc/deg) (n=90 single units, 5 mice) (Fig 2b). The average ratio of photopic CoM to scotopic CoM for single units was 1.07 +/-.45 for spatial frequency (Fig 2c). Temporal frequency tuning was also consistent across light levels (scotopic CoM mean= 2.11 +/-1.08 cyc/sec, photopic CoM mean= 2.46 +/-1.65 cyc/sec) (n=87 single units, 5 mice) (Fig 2e). The average ratio of photopic CoM to scotopic CoM for single units was 1.24 +/-.56 for temporal frequency (Fig 2f). In sum, we find that the CoM of the spatiotemporal tuning properties of V1 single units are consistent across light levels, in keeping with the consistency of responses to movies across light levels. V1 spatiotemporal tuning properties are remarkably invariant to light adaptation state, despite the known changes in the output of the retina (Barlow et al., 1957; Enroth-Cugell and Shapley, 1973).

**Fig 2.**
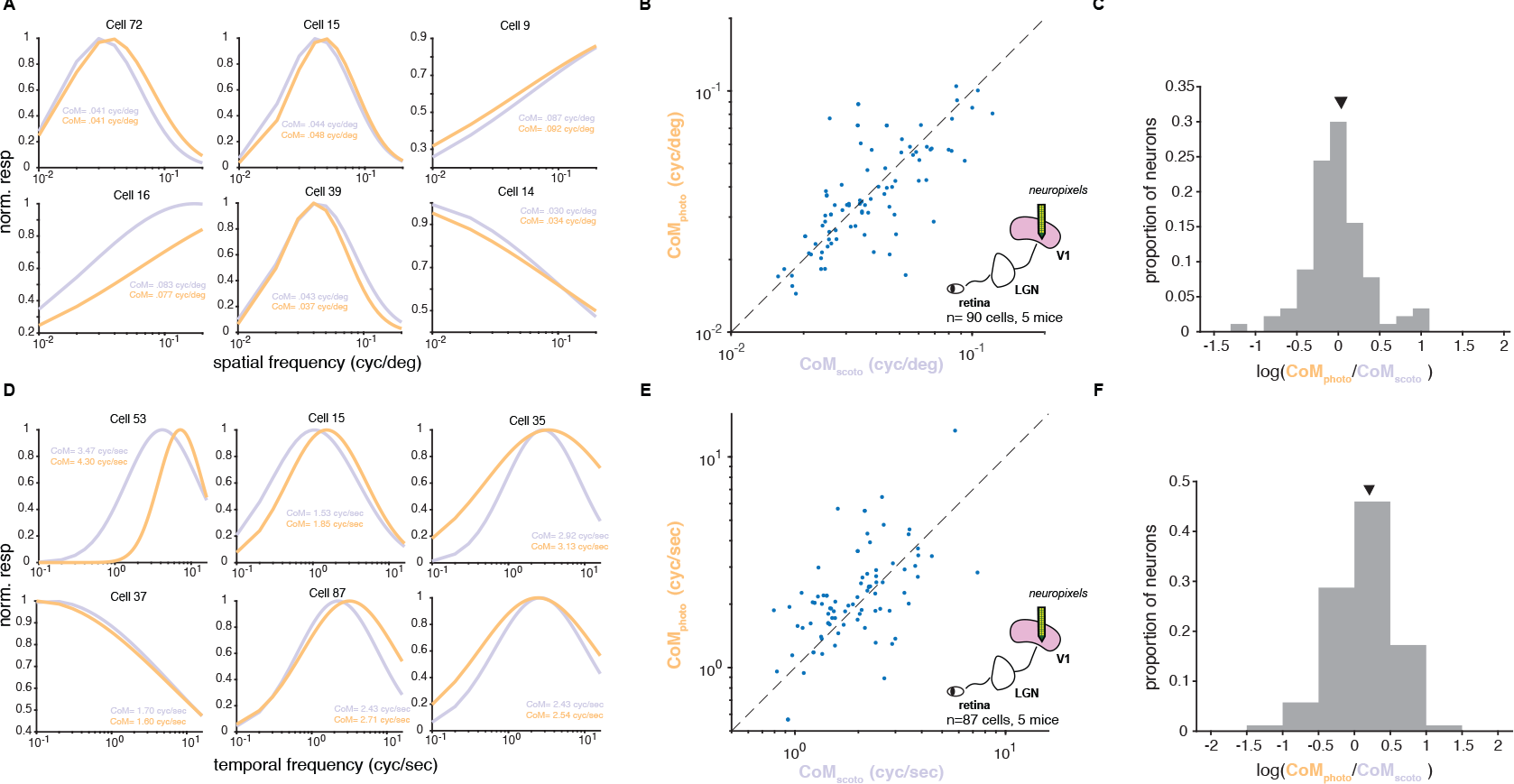
V1 spatial and temporal frequency tuning in scotopic and photopic conditions: A.Example Gaussian spatial frequency tuning curves fit to V1 single unit responses under scotopic (purple) and photopic (orange) conditions. Inset gives center of mass CoM for spatial frequency tuning curves. B. Scatter plot of spatial frequency tuning CoM (cyc/deg) for single units tracked across scotopic and photopic conditions. Dashed line indicates unity. C. Histogram of the log ratio of spatial frequency tuning curve CoM_photo_ /CoM_scoto_ for single units tracked across scotopic and photopic conditions. Arrow indicates mean of distribution. D. Example Gaussian temporal frequency tuning curves fit to V1 single unit responses under scotopic (purple) and photopic (orange) conditions. Inset gives center of mass CoM for temporal frequency tuning curves. E. Scatter plot of temporal frequency tuning CoM (cyc/sec) for single units tracked across scotopic and photopic conditions. Dashed line indicates unity. F. Histogram of the log ratio of temporal frequency tuning curve CoM_photo_ /CoM_scoto_ for single units tracked across scotopic and photopic conditions. Arrow indicates mean of distribution.

### V1 responses have equal variance in scotopic and photopic conditions

Shifts in light level between scotopic and photopic conditions cause changes in the co-variability of RGCs (Greschner et al., 2011; Ruda et al., 2020). For a downstream neuron receiving convergent input from a set of noisy RGCs, the variance of the pooled response will depend on the degree to which RGC responses vary together. In a regime of zero noise correlation, the random fluctuations in RGC activity will tend to cancel out, resulting in a low-variance pooled response. Large, positive noise correlations indicate that RGC responses tend to fluctuate together, which will generate pooled responses with large fluctuations and higher variability. This suggests that the increased covariance of RGC activity in the scotopic regime should lead to greater variance in downstream areas than under the photopic regime.

Surprisingly, the variance of the evoked response to natural movies was significantly larger in the photopic than scotopic condition for single cells (scotopic median=.015 (ΔF/F)^2^, photopic median=.020 (ΔF/F)^2^, p<.001 one-sided Wilcoxon rank sum test) (Fig. 3a, left panel). This is inconsistent with our expectation based on increases in the RGC noise correlations in scotopic conditions. One reason the variability may increase is due to a shift in the mean response magnitude, independent of the light condition. Given the well-known Poisson-like relationship between mean and variance of spiking responses in sensory cortex, we asked whether the larger response magnitude in the photopic case could explain the larger variance (Tomko and Crapper, 1974, Dean 1981). We therefore examined the variance as a function of mean response magnitude and found that the mean-variance relationship is constant between light conditions (Fig. 3a) (two sample t-test, p>.05 in all bins Wilcoxon rank sum test, scotopic slope = .224, photopic slope= .200). We also found a constant mean-variance relationship in V1 responses to gratings. While variance of evoked responses to gratings was significantly larger in the photopic than scotopic condition (scotopic median=6.68 x 10^-4^ (ΔF/F)^2^, photopic median=8.53 x 10^-4^ (ΔF/F)^2^, p<.001 one-sided Wilcoxon rank sum test), variance binned as a function of mean response magnitude was constant (two sample t-test, p>.05 in all bins Wilcoxon rank sum test, scotopic slope = .013, photopic slope= .016) (Supplementary Fig. 1f). The constant relationship between mean and variance of neural responses across light conditions suggests that the upstream changes in noise correlation have little impact on the mean-variance relationship of V1 responses.

**Fig 3.**
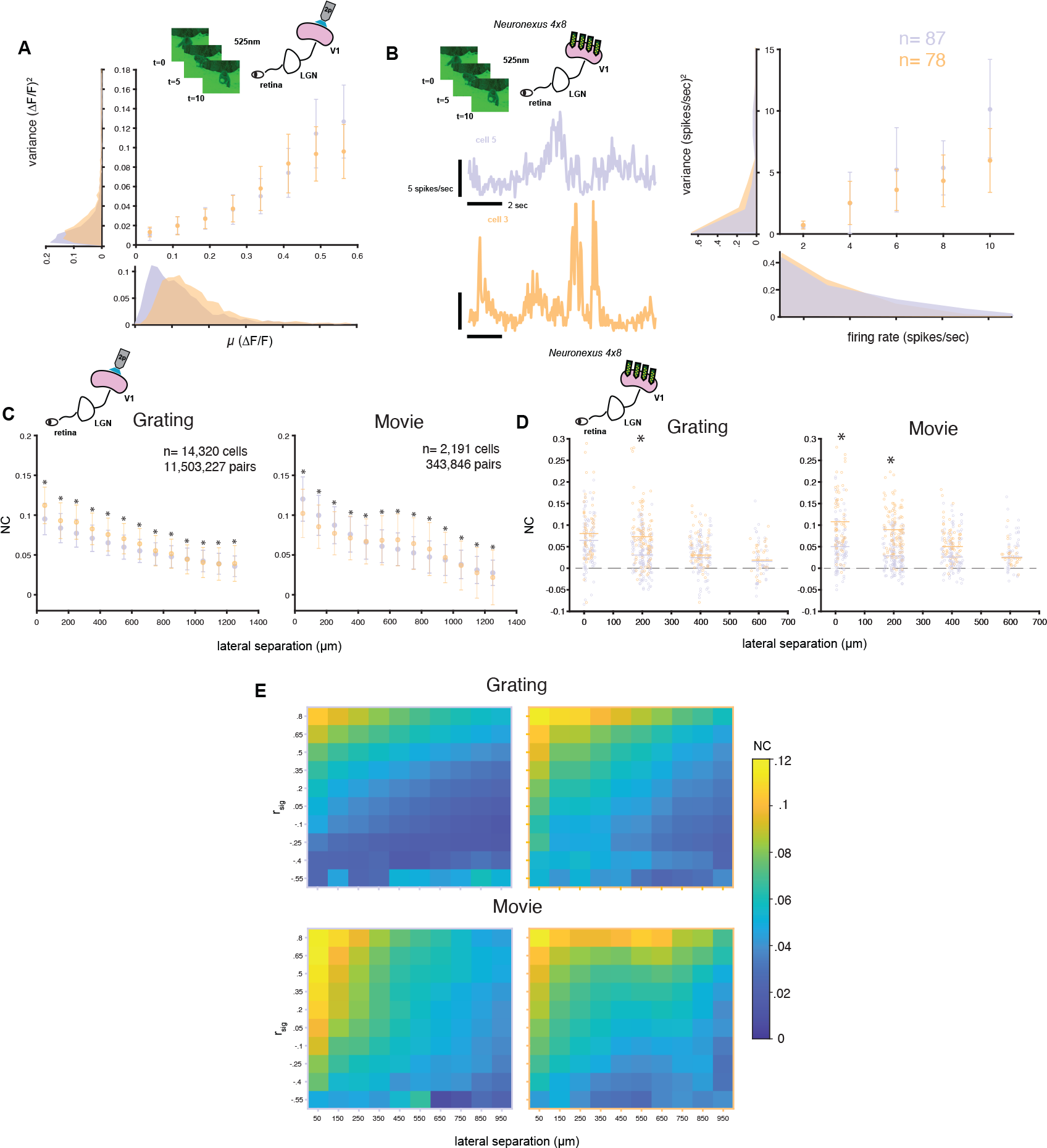
V1 noise correlations in scotopic and photopic conditions: A. Movie response variance plotted as a function of response mean (╭) for all cells in scotopic (purple) and photopic (orange) conditions, recorded with 2-photon calcium imaging. Variance ((′F/F)^2^) binned by ╭ in 8 equally sized bins spanning 0 to .6 F/F. Scatter points indicate average variance within each bin, bars indicate +/-.5 standard deviation. Marginal distributions of mean (bottom) and variance (left) for all cells in in scotopic (purple) and photopic (orange) conditions. B. (Top Left) Experimental design for electrophysiology recordings. Awake mice were head fixed on a treadmill with a 4x8 Neuronexus probe implanted in V1 while viewing 10-second natural movies presented under scotopic and photopic conditions. (Bottom left) Example V1 single unit trial-averaged spiking responses to one movie at scotopic (purple) and photopic (orange) conditions with firing rates computed for 50ms bins. (Right) Movie response variance plotted as a function of mean firing rate (spikes/sec) for all cells in scotopic (purple) and photopic (orange) conditions. Variance ((spikes/sec)^2^) binned by mean in 5 equally sized bins spanning 2 to 10 spikes/sec. Scatter points indicate average variance within each bin, bars indicate +/-.5 standard deviation. Marginal distributions of mean (bottom) and variance (left) for all single units in in scotopic (purple) and photopic (orange) conditions. C. Noise correlations between all pairs of simultaneously imaged V1 cells in scotopic (purple) and photopic (orange) conditions computed for both grating and movie stimulus sets plotted as a function of lateral distance between the pair, recorded with 2-photon calcium imaging. Data grouped by lateral separation into bins spanning 50 ╭m. Scatter points indicated mean noise correlation in each bin, and bars indicate +/-.5 standard deviation. Asterisks indicate p<.05 for Wilcoxon rank sum test. D. Noise correlations between all pairs of simultaneously recorded V1 single units in scotopic (purple) and photopic (orange) conditions computed for both grating and movie stimulus sets plotted as a function of lateral distance between the pair. Horizontal bars indicate the mean noise correlation within each distance bin. Asterisks indicate p<.05 for Wilcoxon rank sum test. E. Heat maps of noise correlations binned by the signal correlation (r_sig_) and lateral separation of each imaged cell pair in scotopic (purple border) and photopic (orange border) conditions for both grating and movie stimulus sets. Value in each square is the mean noise correlation for cell pairs with the corresponding r_sig_ and lateral separation. Color axis is uniform across all plots.

The fluorescent signal which we take as the neural response in these analyses represents the spiking activity of individual cells convolved with a temporally slow, and noisy, calcium kernel. As a result, small changes in the spiking rate of a V1 neuron may not be detectable in the fluorescent response, masking changes in the variance that may exist in V1 spiking response across light conditions. We therefore recorded the spiking response of V1 cells using a four-shank multi-electrode array during repeated presentation of movies and gratings at scotopic and photopic luminance levels. Due to difficulty in tracking the same single units in an awake mouse across the duration of these experiments, we did not attempt to match units across light conditions.

To assess the mean-variance relationship between light conditions, we binned single units based on mean evoked firing rate. As with our imaging findings, there were no significant differences in response variance as a function of mean response magnitude between luminance conditions for natural movies or static gratings (Fig. 3b, Supplemental Fig 1g) (two sample t-test, p>.05 in all bins, Wilcoxon rank sum test). Neither calcium signals nor spiking data from V1 exhibited the expected shift in variance that should accompany a change in retinal noise correlations.

### V1 noise correlations are invariant to light adaptation state

Records from the retina demonstrate that noise correlations increase substantially in the scotopic state and that a specific downstream population decoder would fail if this change in noise structure is not taken into account (Greschner et al., 2011; Ruda et al., 2020). Such changes in the magnitude of retinal noise correlations could generate a commensurate change in the magnitude of V1 noise correlations. To examine whether the noise correlation signature of the scotopic state propagates into V1, we computed the noise correlations for all cortical neuron pairs (NC_V1_). We computed NC_V1_ using the single trial responses from both our natural movie and grating imaging datasets. We find no signature of excess noise in the scotopic state in V1.

Any change in the noise structure from the retina would be most apparent for the closest V1 cell pairs, which share retinotopic inputs. Even for the closest cell pairs, separated by less than 50 ╭m, the distributions of NC_V1_ for the scotopic and photopic states overlap, but we did uncover significant shifts in noise correlations between scotopic and photopic conditions. For gratings, the mean NC_V1_ for neighboring cells declined from 0.129 +/-.048 to 0.104 +/-.041 as light level shifted from photopic to scotopic levels. This is surprising given our expectation that NC_V1_ would increase under scotopic conditions along with retinal noise correlations (Ruda et al., 2020). For the natural movies we observed a shift in the opposite direction amongst neighboring cell pairs: photopic NC_V1_ = 0.118+/-.067 and scotopic NC_V1_ = 0.136 +/-.059. These NC_V1_ distributions are highly overlapping, exhibit mild shifts in both directions, and are statistically significant because of a large number of cell pairs examined (n=29309 pairs for gratings, n= 5606 pairs movie). These NC_V1_ distributions do not reveal a systematic change in noise correlation magnitude with light level as was found for neighboring RGCs.

We next examined how NC_V1_ depended on the lateral distance between cell pairs. As for RGCs, NC_V1_ systematically declines with distance (Fig. 3c), though any differences between scotopic and photopic conditions remain mild and inconsistent for grating and movie responses. These differences across light levels are often statistically significant because of the massive number of cell pairs from which we recorded (n=1.15 x 10^7 pairs for gratings, n= 3.44 x 10^5 pairs for movie). Knowledge of light condition explains little of the variance in the NC_V1_ data for cell pairs separated by a given distance. If NC_V1_ distributions across light levels resembled the data from the retina, with distributions separable and tightly grouped around the mean for each light condition, then the variance explained by knowledge of light condition should be substantial (Ruda et al., 2020). Light condition in V1, however, accounted for only a small percentage of the variance in NC_V1_ data, reflecting both the strong overlap across light conditions and high within-condition variability in NC_V1_ for neighboring cell pairs (movie var explained=2.3%, grating var explained= 6.9%).

We also noted that NC_V1_ exhibited a high degree of variability within a given spatial distance (std = .04-.05) and light condition, and this variability was constant across cortical space. Some of this variability may be related to difference in the functional properties of cell pairs. Previous studies have found that pairwise signal correlations (r_sig_) are predictive of NC_V1_ (Smith and Kohn, 2008; Ko et al., 2011). Indeed, there is a strong dependence of NC_V1_ on r_sig_ across cortical space in our dataset (Fig. 3e). By binning over cortical distance and r_sig_, we find that most of the variability in NC_V1_ magnitude is explained by a product of two functions defining the interaction of a linear decay over cortical distance and an exponential decay with r_sig_ (Smith and Kohn, 2008).

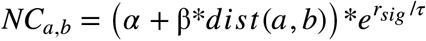

Fitting this 3-parameter model to the binned NC_V1_ data reveals that this relationship remains constant between light conditions (Table 1).

**Table 1:** Model fits for NC_V1_ as a function of lateral cortical distance and r_sig_.

|  | Scotopic | Photopic | Combined |
| --- | --- | --- | --- |
| **Grating** |  |  |  |
| $\alpha$ | .105 | .121 | .110 |
| $\beta$ | .057 | .074 | .064 |
| $\tau$ | 1.06 | 1.47 | 1.37 |
| Variance Explained | 77.2% | 92.5% | 87.9% |
| **Movie** |  |  |  |
| $\alpha$ | .121 | .119 | .119 |
| $\beta$ | .082 | .047 | .065 |
| $\tau$ | 2.34 | 1.51 | 1.82 |
| Variance Explained | 88.0% | 90.2% | 92.9% |

Differences in parameters for our model fit to NC_V1_ data from each light condition could reflect meaningful differences in the relationship between NC_V1_, r_sig_, and cortical distance, or simply be due to overfitting. To assess whether the model parameters differ significantly between light conditions, we employed the Akaike information criterion (AIC, see METHODS) (Akaike, 1974). We compared two candidate models, one in which NC_V1_ was estimated by separate α, β, and τ parameters for each light condition (6 total free parameters), and one in which NC_v1_ was estimated with the same 3 parameters regardless of light condition (combined model). For model fits to both grating and movie stimuli, we find that the AIC is lower for 3-parameter model, indicating that NC_v1_ is well estimated by a function of r_sig_ and cortical distance that is invariant to light level (Grating AIC: 5.22 (6 parameter) vs 3.23 (3 parameter), Movie AIC: 6.05 (6 parameter) vs 3.30(3 parameter)). In sum, our calcium imaging dataset reveals that light adaptation state is not a determinant of NC_V1_.

As discussed above, the coarse temporal resolution of calcium imaging may mask an effect of light adaptation state that is apparent in the spiking response. To address this concern, we also computed NC_V1_ for single unit responses to identical grating and natural movie stimulus sets at scotopic and photopic luminance levels.

Using a multi-electrode array, we were able to map NC_V1_ across ∼650 ╭m of cortex in a manner analogous to our imaging experiments. We again found that NC_V1_ distributions were highly overlapping between light conditions. For pairs of V1 single units recorded on the same electrode shank, and therefore separated by less than ∼40 ╭m laterally, NC_V1_ declined on average from photopic to scotopic conditions for the responses to both gratings (photopic NC_V1_ =.088 +/-.067, scotopic NC_V1_ =.066 +/-.045) and movies (photopic NC_V1_ =.111 +/-.070, scotopic NC_V1_ =.047 +/-.042). Across lateral cortical space, photopic NC_V1_ was higher on average than scotopic NC_V1_, in stark contrast with prior results in the retina (Fig. 3d). Given the high degree of overlap in the NC_V1_ distributions, light condition accounted for little of the variance in our NC_V1_ spiking data (movie var explained=1.75%, grating var explained=.75%).

In sum, our data reveal that there is not systematic shift in NC_v1_ between scotopic and photopic light conditions, as measured with both 2-photon imaging and electrophysiology. The clearest difference between our electrophysiology and imaging results is that the overall NC_V1_ was smaller on average for the spiking response. This difference may be due to some neuropil contamination in single cell response traces extracted from calcium imaging data, particularly for the transgenic mice used in this study, which express dense fluorescent labeling of excitatory cells in V1 (see Methods, Pachitariu et al., 2017).

### Invariant encoding of orientation in V1 across light adaptation states

Given the consistency of the mean, variance, and covariance of V1 responses between light conditions, we hypothesized that the population-level representation of visual orientation between light conditions should remain constant. The discriminability of the V1 population responses for a pair of orientations is a function of the difference in the average response across all cells (d╭) and the covariance of the responses across trials (Σ). Due to limited experimental repeats of each stimulus, estimates of Σ for large neural populations tend to be unreliable.

Instead, the high dimensional neural population data can be projected into a lower dimensional space from which d′ may be estimated. Heller and David developed such a method in which neural data are projected into a 2-dimensional space (dDR space) defined by d╭ the axis that maximizes the differences in the mean responses) and the orthogonal component of the first eigenvector of the covariance matrix (e_1,_ the “noise axis”). e_1_ and d╭ are not necessarily orthogonal. If e_1_ is orthogonal to d╭ then d-prime can be computed directly from d╭. If, however, e_1_ is not orthogonal, then the maximal discrimination axis is tilted in this 2-dimensional space (Fig. 4a) (Heller and David, 2022).

**Fig 4.**
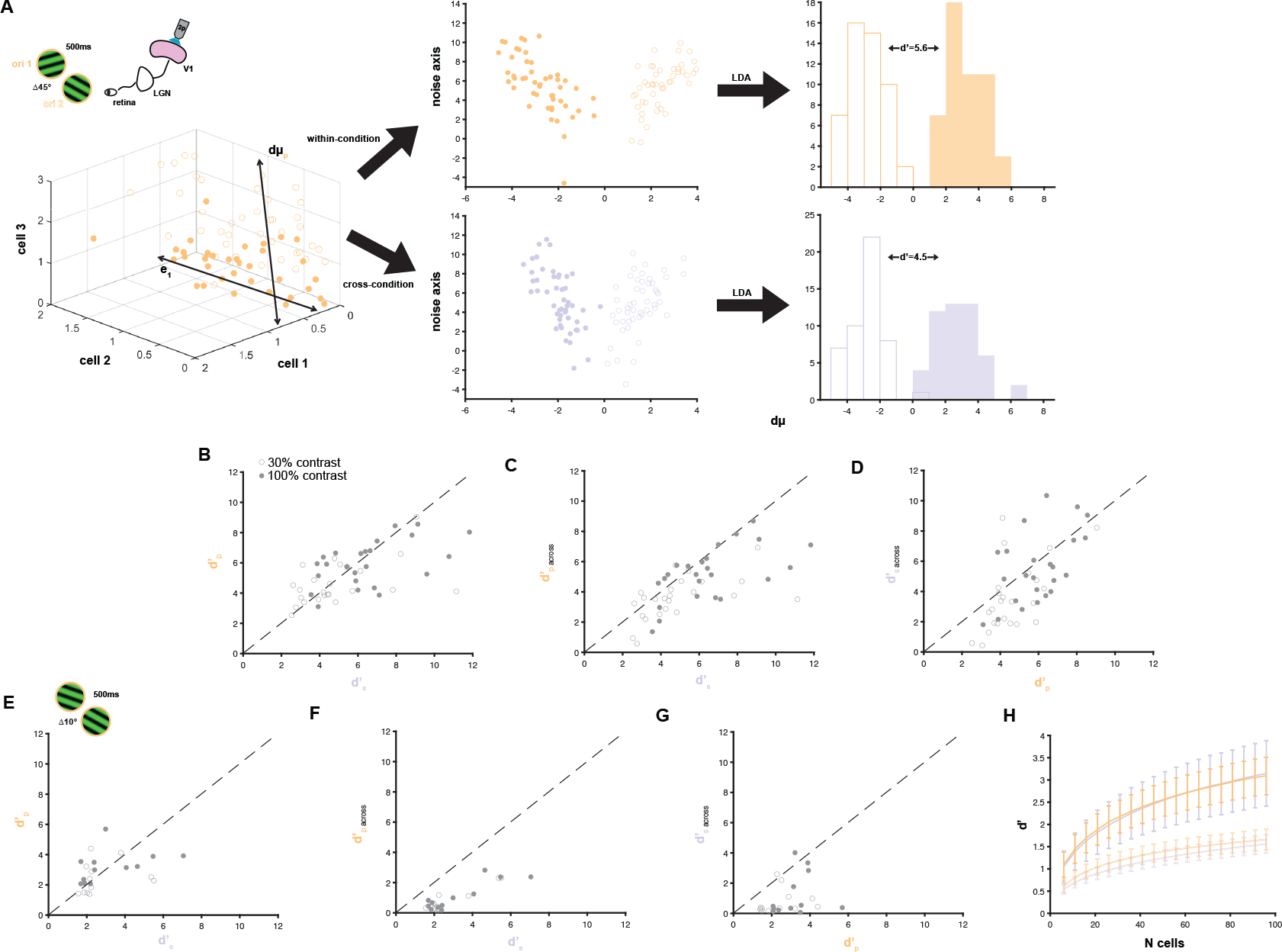
Discriminability of the V1 population code for orientation under scotopic and photopic conditions: A.Example of dDR dimensionality reduction method for orientation discrimination on imaging data from V1. (Top Left) Experimental design. Awake mice were head fixed on a treadmill underneath a two-photon microscope while viewing 50 repeats of static gratings at 4 orientations separated by 45° at 30% and 100% contrast under scotopic (purple) and photopic (orange) conditions. To compare V1 responses in these two regimes, the same V1 neurons were tracked across conditions. (Bottom left) Example of V1 population responses plotted for 50 trials of gratings at 2 orientations. Each axis corresponds to the single trial evoked response (Δ F/F) of one V1 cell. Filled scatter points indicate the population response to orientation 1, open scatter points indicate the population response to orientation 2. d╭ is the axis in neural space that maximally separates the mean response of the population to each stimulus. e_1_ is the first eigenvector of the covariance matrix for the neural data. d╭ and e_1_ are computed for each pair of stimuli. (Top Middle) V1 population responses are projected into the dDR space-a 2 dimensional space defined by d╭ and the component of e_1_ perpendicular to d╭ (the “noise axis”). This operation is performed on data from the same light condition for which the dDR space was computed (“within-condition”) and (Bottom Middle) on data for the same pair of gratings in the other light condition (“cross-condition”). (Right) To compute the discriminability index for a pair of stimuli (*d′*), we perform linear discriminant analysis (LDA) to find the axis for which responses to the 2 stimuli in the dDR space are maximally separated. B. Scatter plot of the discriminability index of Δ45° computed for each pair of stimuli in each experiment independently for scotopic (d′_s_) and photopic (d′_p_) conditions using the dDR method. C. Scatter plot of the discriminability index of Δ45° computed for each pair of stimuli in each experiment using scotopic data to fit the dDR space into which both the scotopic and photopic data were projected to compute d′_p_ and d′_s, across_. D. Scatter plot of the discriminability index of Δ45° computed for each pair of stimuli in each experiment using photopic data to fit the dDR space into which both the scotopic and photopic data were projected to compute d′_s_ and d′_p, across_. E. Scatter plot of the discriminability index of 10° computed for each pair of stimuli in each experiment independently for scotopic (d′_s_) and photopic (d′_p_) conditions using the dDR method. F. Scatter plot of the discriminability index of 10° computed for each pair of stimuli in each experiment using scotopic data to fit the dDR space into which both the scotopic and photopic data were projected to compute d′_s_ and d′_p, across_. G. Scatter plot of the discriminability index of 10° computed for each pair of stimuli in each experiment using photopic data to fit the dDR space into which both the scotopic and photopic data were projected to compute d′_p_ and d′_s, across_. H. Discriminability index of 10° at 30% contrast as a function of random sample size of simultaneously recorded cells. Scatter points indicated mean d′ for each cell sample size, and bars indicate +/-.5 standard deviation.

In this 2-D space, the discriminability index for a pair of stimuli (*d′*) is

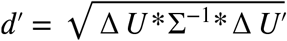

Where *U* is a 2D vector of the difference in mean response to the two conditions, and Σ is the 2x2 covariance matrix.

To compare the discriminability of the V1 code across illumination states, we used calcium imaging to track V1 cells during presentation of static gratings separated by 45° at scotopic and photopic light levels (n=14,320 V1 cells, 1193.3 +/-734.4 cells/session, Supplementary Figure 2a, b). We find that *d′* for 45° discrimination in the V1 population is well matched between scotopic and photopic conditions at both 30% and 100% contrasts (scotopic mean 5.72 +/-2.34, photopic mean 5.30 +/-1.59)(Fig 4b). Further, the choice of dimensionality reduction method does not qualitatively impact this result, as employing the partial least squares method yields similar measurements of *d′* as the dDR method (scotopic mean 2.12 +/-1.84, photopic mean 2.79 +/-1.82)(Supplementary Fig 2d) (Rumyantsev et al., 2020).

This analysis indicates that V1 encodes coarse orientation differences equally well for both light levels, but not whether the structure of the code is constant. To determine if the structure of the V1 code for orientation is constant between light levels, we examined how *d′* changes if the population responses are projected into the dDR space computed for the other light level. If the structure of the population code changes significantly across light conditions, then the dDR space will not generalize to data from across conditions, and the *d′* computed for cross-condition data will be much smaller than for within-condition data. Instead discriminability is largely conserved in the dDR space across light conditions (scotopic across mean= 4.58 +/-2.48, photopic across mean=4.40 +/-1.77, Fig 4c, d).

The coarse discrimination of 45° differences in grating orientation is not challenging for the V1 population, as evident in the large *d′* values which we report. Such coarse discrimination may not be sensitive to changes in the retinal output due to the transition between rod and cone mediated vision. We therefore employed a finer discrimination task, where response distributions overlap across stimuli. To examine whether V1 encoding of small orientation differences is invariant to light level, we presented static gratings separated by 10° (7,472 cells, 6 experimental sessions, 1245 +/-823 cells/session). We again found that the discriminability of the V1 population code does not change significantly between light conditions (scotopic mean= 2.98 +/-1.54, photopic mean=2.83+/-1.09) (Fig. 4e). The dDR space did not generalize as well for cross-condition data as it had for 45° discrimination, as *d′* computed for cross-condition data was much smaller than for within-condition data (scotopic across mean=.989+/-1.15, photopic across mean=.944+/-.849, Fig. 4f, g). Adding more dimensions to our reduced dimensional space does not increase within-condition *d′* values (scotopic mean=2.07+/-1.83, photopic mean=2.83+/-1.84), but does improve the generalization of orientation discrimination for cross-condition data (scotopic across mean=1.36+/-.986, photopic across mean=1.34+/-.910, see METHODS) (Supplementary Fig. 2e, f).

Discriminability at 10° orientation differences was significantly lower than 45° differences, but d′ remains high when data from large V1 populations are included. To determine whether discriminability may be more strongly limited by light level when only a small neural population is considered, we randomly subsampled our neural populations and measured *d′* . We find a consistent relationship between population size and *d′* between light conditions for both 45° and 10° discrimination at 30% contrast, in which *d′* grows with population size (Fig. 4h). In sum, the discriminability for orientation in the V1 code is consistent across light levels, even in the limit of fine orientation differences and small neural populations.

The dramatic change in retinal noise correlations observed across light conditions could not only alter the projections into the dDR space, but could alter the primary axes along which the neural population covaries (e1). To test whether there is a change in the V1 noise structure we computed e_1_ separately for scotopic (e_1s_) and photopic (e_1p_) conditions for every stimulus pair. The dot product of e_1s_ and e_1p_ quantifies the alignment of these two axes between light conditions. Across all neural populations and stimulus pairs in our dataset, the mean cross-condition dot product was high, indicating that V1 responses co-vary along the same primary axis in scotopic and photopic conditions (mean=.89 +/-.09, Supplementary Fig. 2c).

### LGN receptive field properties shift with light adaptation state

We have shown that, despite the known changes in the output of the retina with luminance, the spatial and temporal frequency tuning, as well as the variance and covariance of the V1 population does not shift substantially between the scotopic and photopic states. It is unclear from these results alone why the functional shifts in RGC activity are not detectable in cortex. One possibility is that the changes in RGC firing do not significantly impact responses immediately downstream in the lateral geniculate nucleus (LGN). Alternatively, the LGN could inherit functional shifts with light level, but these shifts do not persist downstream in V1. To adjudicate between these two hypotheses, we densely sampled LGN single unit activity using the Neuropixels 1.0 probe while awake mice viewed stimuli under scotopic and photopic conditions.

We first asked whether the receptive field changes observed in RGC activity across light levels was also apparent in LGN responses. We used the same set of drifting grating stimuli which we employed in our V1 experiments to measure the spatial and temporal tuning of LGN single units. The selectivity of the LGN population shifted significantly toward higher spatial frequencies in the photopic state, consistent with observations that spatial frequency tuning shifts from low-pass to band-pass as light levels increase (scotopic CoM mean= .041 +/-.025 cyc/deg, photopic CoM mean= .054 +/-.033 cyc/deg) (p<.001 one-sided Wilcoxon rank sum test) (n=124 single units, 6 mice) (Fig 5b) (Barlow et al., 1957). The average ratio of photopic CoM to scotopic CoM for single units is 1.48 +/-.99 for spatial frequency (Fig 5c). There was a trend for tuning to shift to higher temporal frequencies in the photopic relative to scotopic state, but this increase was not statistically significant (scotopic CoM mean= 3.10 +/-2.38 cyc/sec, photopic CoM mean= 3.44 +/-2.38 cyc/deg) (p=.1 one-sided Wilcoxon rank sum test) (n=77 single units, 6 mice) (Fig 5e). The average ratio of photopic CoM to scotopic CoM for single units is 1.35 +/-.97 for temporal frequency (Fig 5f).

**Fig 5.**
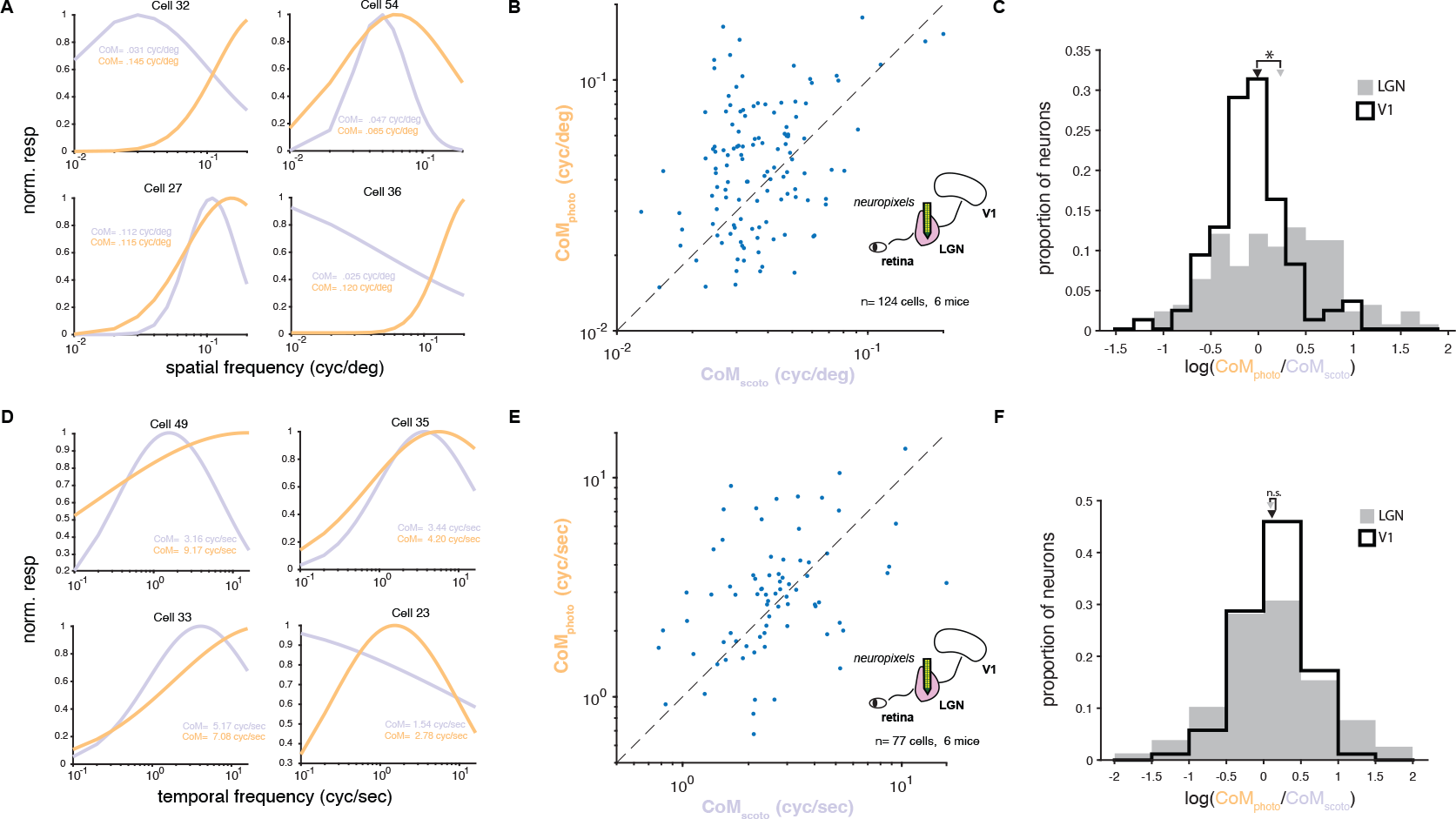
LGN spatial and temporal frequency tuning in scotopic and photopic conditions: A. Example Gaussian spatial frequency tuning curves fit to LGN single unit responses under scotopic (purple) and photopic (orange) conditions. Inset gives center of mass CoM for spatial frequency tuning curves. B. Scatter plot of spatial frequency tuning CoM (cyc/deg) for LGN single units tracked across scotopic and photopic conditions. Dashed line indicates unity. C. Histogram of the log ratio of spatial frequency tuning curve CoM_photo_ /CoM_scoto_ for LGN and V1 single units tracked across scotopic and photopic conditions. Arrows indicates means of distributions. Asterisk indicates p<.05 for one-sided Wilcoxon rank sum test between LGN and V1 distributions. E. Example Gaussian temporal frequency tuning curves fit to LGN single unit responses under scotopic (purple) and photopic (orange) conditions. Inset gives center of mass CoM for temporal frequency tuning curves. F. Scatter plot of temporal frequency tuning CoM (cyc/sec) for LGN single units tracked across scotopic and photopic conditions. Dashed line indicates unity. G. Histogram of the log ratio of temporal frequency tuning curve CoM_photo_ /CoM_scoto_ for LGN and V1 single units tracked across scotopic and photopic conditions. Arrows indicate means of distributions.

These LGN records are distinct from our V1 records as the magnitude of shifts in spatial frequency tuning with light level are much larger for LGN than V1 single units (LGN CoM ratio mean=1.48 +/-.99, V1 CoM ratio mean=1.07 +/-.45, p<.001 one-sided Wilcoxon rank sum test)(Fig 5c), while the magnitude of temporal frequency tuning shifts are similar in LGN and V1 (LGN CoM ratio mean=1.35 +/-.97, V1 CoM ratio mean= 1.24 +/-.56, p=.59 one-sided Wilcoxon rank sum test) (Fig 5f). Therefore, the LGN shifts spatial frequency tuning in the same direction as RGCs with light level while V1 neurons retain their spatial frequency tuning. Interestingly, changing illumination had little effect on temporal frequency tuning in either the LGN or V1.

### Variance and correlations vary with light level in the LGN

We also sought to determine whether luminance alters the variance and covariance of LGN neurons as it does RGCs. To do so, we measured single unit activity along the dorso-ventral extent of the LGN to repeated presentations of grating stimuli under scotopic and photopic conditions in order to compute the variance and covariance of the responses (Fig 6a). LGN spiking responses are significantly more variable as a function of mean firing rate in the scotopic than photopic regime (Fig 6b) (p<.05 Wilcoxon rank sum test, scotopic slope= 4.8, photopic slope=3.1). This increased variance in the scotopic state is consistent with the notion that the variance of a neuron’s response will increase when pooling from a more highly correlated set of inputs.

**Fig 6.**
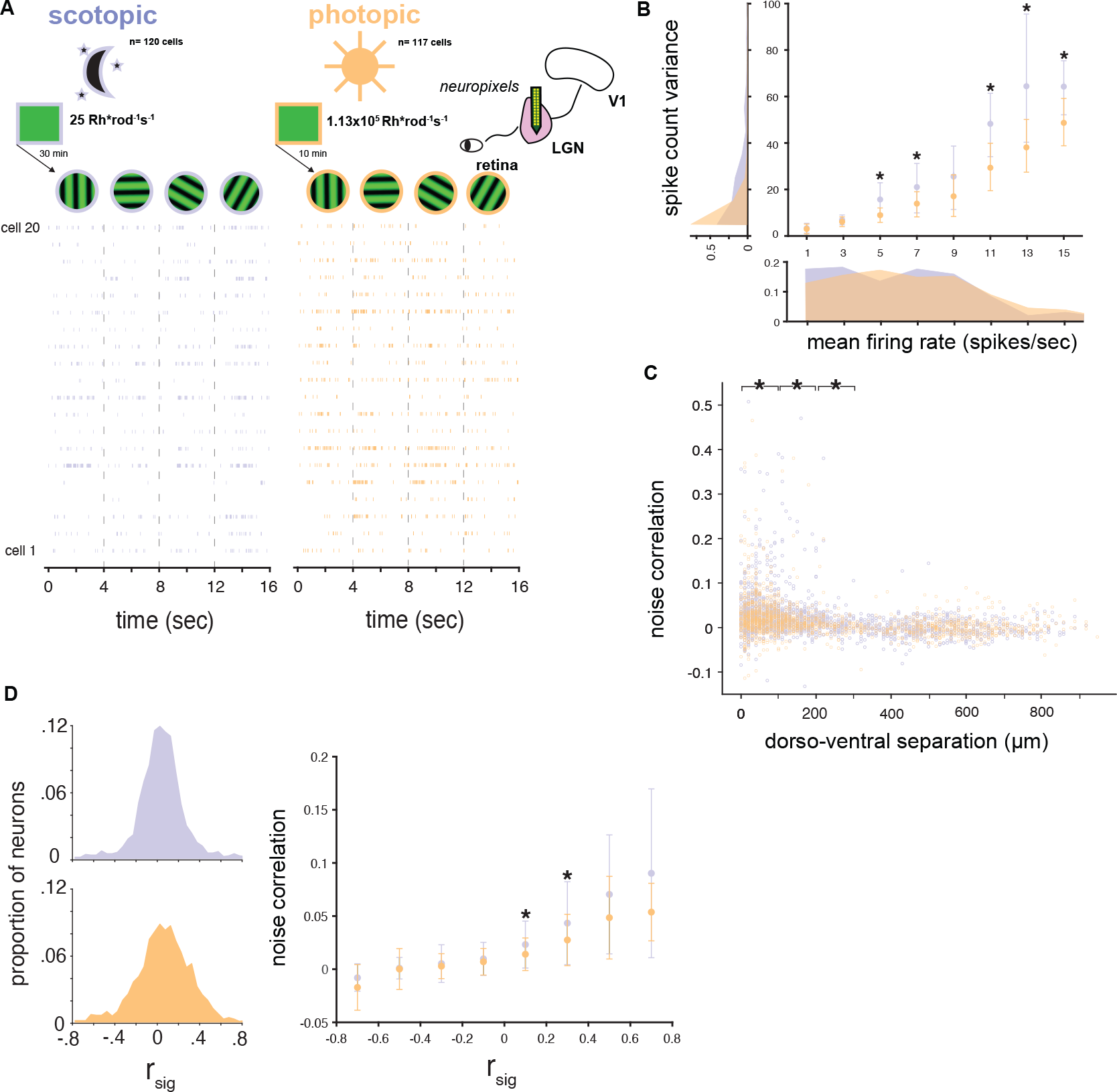
LGN noise correlations in scotopic and photopic conditions: A. (Top) Experimental design. Awake mice were head fixed on a treadmill with the Neuropixels 1.0 probe implanted in LGN while viewing 100 repeats of 2 sec long drifting gratings at 4 directions under scotopic (purple) and photopic (orange) conditions. (Bottom) Example single trial spiking response for 20 simultaneously recorded LGN single units to 4 drifting gratings under scotopic and photopic conditions. B. Response variance plotted as a function of mean firing rate (spikes/sec) for all cells in scotopic (purple) and photopic (orange) conditions. Variance ((spikes/sec)^2^) binned by mean in 5 equally sized bins spanning 1 to 5 spikes/sec. Scatter points indicate average variance within each bin, bars indicate +/-.5 standard deviation. Marginal distributions of mean (bottom) and variance (left) for all single units in in scotopic (purple) and photopic (orange) conditions. Asterisk indicates p<.05 for one-sided Wilcoxon rank sum test. C. Noise correlations between all pairs of simultaneously recorded LGN single units in scotopic (purple) and photopic (orange) conditions plotted as a function of the dorso-ventral distance between the pair. Asterisks indicate significantly larger scotopic noise correlations in bins of 100 ╭m (p<.05 one-sided Wilcoxon rank sum test). D.(Left) Histogram of signal correlation (r_sig_) between all simultaneously recorded geniculate cell pairs computed separately for scotopic (purple) and photopic (orange) conditions. (Right) Noise correlation plotted as a function of r_sig_ for each cell pair, binned by r_sig_ in 8 equally sized bins spanning -.75 to .75. Scatter points indicate average noise correlation within each bin, bars indicate +/-.5 standard deviation. Asterisk indicate significantly larger scotopic noise correlations in a bin (p<.05 one-sided Wilcoxon rank sum test).

Noise correlations between LGN pairs also increase in scotopic conditions. Prior work has shown that large, positive correlations exist primarily between RGCs of the same ON/OFF polarity and type, with small retinotopic offsets (Ala-Laurila et al. 2011; Greschner et al., 2011; Ruda et al. 2020). If retinal noise correlations are inherited by the LGN population, then we expect to observe large, positive correlations between nearby geniculate pairs (NC_LGN_) receiving input from retinotopically overlapping RGCs of the same polarity and type; these pairs will also have high r_sig._ We therefore examined cell pairs separated by less than 300 ╭m and found that NC_LGN_ is significantly higher in the scotopic than photopic state (scotopic mean = .032 +/-. 061, photopic mean= .024 +/-.049, p<.01 Wilcoxon rank sum test) (Fig. 6c). The increase in NC_LGN_ in scotopic luminance is due to an increase in NC_LGN_ for cell pairs with high r_sig_, consistent with the prediction that changes in NC_LGN_ across light conditions are due to neighboring geniculate pairs receiving input from the same RGC class (Fig. 6d).

Though there is only a modest increase in NC_LGN_ relative to that observed in the retina under scotopic conditions, it is clearly distinct from the distributions which we observed for NC_V1_, where correlations were equal or larger in the photopic than scotopic conditions. We conclude that this result constitutes a recapitulation of noise distributions observed across retinal space under scotopic and photopic conditions (Ruda et al. 2020). In addition, geniculate pairs with zero or negative r_sig_ have near-zero NC_LGN_ on average, indicating that NC_LGN_ is primarily driven by shared inputs from the retina. This is also apparent in the decline to zero NC_LGN_ for distant cell pairs-which receive retinotopically disparate inputs (Piscopo et al., 2013).

### A model of afferent convergence explains V1 spatial tuning and noise structure across light levels

Our experimental results show that changes in the spatial tuning and noise correlations of the RGC population with light level persist in LGN, but not downstream in V1. We propose a simple physiological mechanism to explain these results based on a convergent feedforward architecture in which parallel processing streams from the retina and LGN are integrated by single cortical cells.

In studying the downstream consequences of functional shifts in the retina, it is important to consider the diverse classes of RGCs present in the rodent retina, which show heterogeneous shifts in their functional properties and correlation structure across light adaptation states (Baden et al., 2016; Ruda et al., 2020; 2022). Since convergence of these functionally diverse peripheral inputs is a canonical substrate of the V1 receptive field, we investigated the effects of pooling from retinal afferents with both ON and OFF polarity and diverse functional class (Hubel and Wiesel, 1962; Ferster et al. 1996; Chung and Ferster, 1998, Lien and Scanziani, 2013). We constructed a two-layer network model in which V1 receptive fields and noisy responses emerge from the pooling of units in the input layer (Ringach, 2007; Kanitcheider et al., 2015, Pattadkal et al. 2018). For simplicity, we treat LGN as a relay of the retinal output. The model is constrained by the regular spacing of RGC receptive fields of a given class, as well as the relative receptive field sizes of RGCs and V1 cells, such that each model V1 cell pools from a sparse set of peripheral inputs (Fig 7a) (Ringach, 2007, 2021).

**Fig 7.**
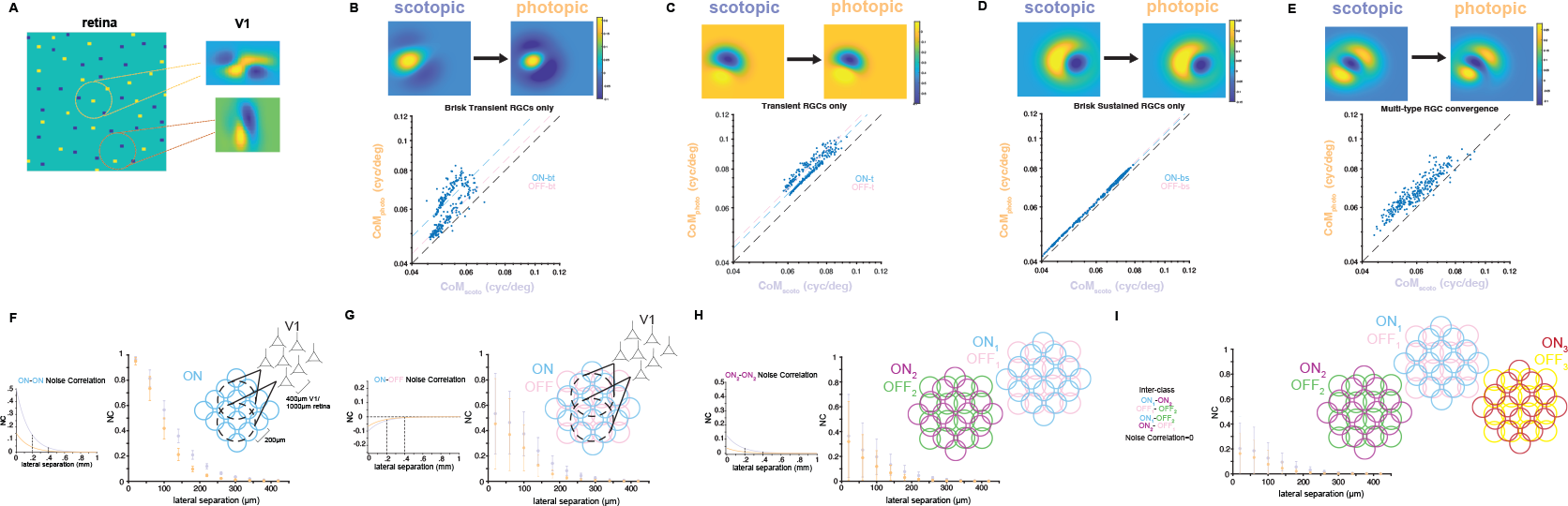
A feedforward model of V1 in scotopic and photopic conditions: A. Schematic of modeling approach for V1 receptive fields. ON (yellow) and OFF (blue) RGCs have circular Gaussian receptive fields and tile retinotopic space in a mosaic pattern. Each V1 unit pools from a local set of RGCs with weights defined by a circular Gaussian in retinotopic space (dashed line). We simulate a dense population of V1 neurons with receptive field centers shifting as a function of lateral separation in cortex. B. Model V1 receptive fields pooling from a single class of RGCs with large modulation of receptive field size across light levels, resembling the functional properties of the Brisk Transient RGC class in rodent retina. (Top) Example receptive field of a model V1 unit in scotopic and photopic conditions. (Bottom) Scatter plot of spatial frequency tuning CoM (cyc/deg) for model V1 units under scotopic and photopic conditions. Black dashed line indicate unity. Colored dashed lines indicate the expected ratio CoM_photo_/CoM_scoto_ for a V1 cell pooling from only ON-bt (blue) or OFF-bt (pink) inputs. C. Model V1 receptive fields pooling from a single class of RGCs with intermediate modulation of receptive field size across light levels, resembling the functional properties of the Transient RGC class in rodent retina. (Top) Example receptive field of a model V1 unit in scotopic and photopic conditions. (Bottom) Scatter plot of spatial frequency tuning CoM (cyc/deg) for model V1 units under scotopic and photopic conditions. Black dashed line indicate unity. Colored dashed lines indicate the expected ratio CoM_photo_/CoM_scoto_ for a V1 cell pooling from only ON-t (blue) or OFF-t (pink) inputs. D. Model V1 receptive fields pooling from a single class of RGCs with minimal modulation of receptive field size across light levels, resembling the functional properties of the Brisk Sustained RGC class in rodent retina. (Top) Example receptive field of a model V1 unit in scotopic and photopic conditions. (Bottom) Scatter plot of spatial frequency tuning CoM (cyc/deg) for model V1 units under scotopic and photopic conditions. Black dashed line indicates unity. Colored dashed lines indicate the expected ratio CoM_photo_/CoM_scoto_ for a V1 cell pooling from only ON-bs (blue) or OFF-bs (pink) inputs. E. Model V1 receptive fields pooling from 3 classes of RGCs, resembling the functional properties of the Brisk Transient, Brisk Sustained, and Transient RGC classes in rodent retina. (Top) Example receptive field of a model V1 unit in scotopic and photopic conditions. (Bottom) Scatter plot of spatial frequency tuning CoM (cyc/deg) for model V1 units under scotopic and photopic conditions. Black dashed line indicates unity. F.(Left) Within-class retinal noise correlations are modeled as a function of lateral separation between each RGC pair with an exponential function. Dashed lines indicate noise correlation magnitude for RGC pairs separated by 200 and 400 ╭m. (Middle) Noise correlations between all model V1 cell pairs plotted as a function of lateral distance between the pair for scotopic (purple) and photopic (orange) conditions. Data grouped by lateral separation into bins spanning 50 ╭m. Scatter points indicated mean noise correlation in each bin, and bars indicate +/-.5 standard deviation. (Right) Schematic of model design. RGCs of a single class and polarity are arranged in a mosaic with 200 ╭m between receptive field centers. V1 neurons pool the responses of a local set of RGCs with weights defined by a circular Gaussian in retinotopic space (dashed line). We simulate a dense population of V1 neurons with receptive field centers shifting as a function of lateral separation in cortex. G. (Left) ON-OFF within-class retinal noise correlations are modeled as a function of lateral separation between each RGC pair with an exponential function. Dashed lines indicate noise correlation magnitude for RGC pairs separated by 200 and 400 ╭m. (Middle) Noise correlations between all model V1 cell pairs as in (E). (Right) Schematic of model design. Two RGC mosaics are arranged in anti-phase, one ON and one OFF. V1 neurons pool the responses of a local set of RGCs with weights defined by a circular Gaussian in retinotopic space (dashed line), and have equal probability of pooling from ON or OFF RGCs. H. Within-class retinal noise correlations for a second class of RGCs are modeled as a function of lateral separation between each RGC pair with an exponential function with differing magnitude and spatial extent from (E, F). (Middle) Noise correlations between all model V1 cell pairs as in (E). (Right) Schematic of model design. 2 sets of ON and OFF RGC mosaics arranged in antiphase. V1 neurons pool the responses of a local set of RGCs with weights defined by a circular Gaussian in retinotopic space (dashed line), and have equal probability of pooling from any RGC. I. (Left) Noise correlations between all model V1 cell pairs as in (E). (Right) Schematic of model design. 3 sets of ON and OFF RGC mosaics arranged in antiphase. V1 neurons pool the responses of a local set of RGCs with weights defined by a circular Gaussian in retinotopic space (dashed line), and have equal probability of pooling from any RGC.

We first use this model to study the impact of changing light level on spatial tuning properties in V1. As light level changes, the connectivity from the periphery to V1 is fixed, but the input receptive fields change according to the data provided in Ruda et al. 2022. Those data show that some RGC classes have receptive field radii are strongly modulated by luminance, such as Brisk Transient or Transient RGCs, which show up to a 20% decrease in receptive field radii between scotopic and photopic conditions. A V1 cell pooling from just these RGCs would show a shift in tuning towards higher spatial frequencies under photopic conditions (Fig 7b, c). Still other RGC classes, such as Brisk Sustained RGCs, have relatively invariant receptive field sizes across light levels. A V1 cell pooling from these RGCs would have invariant spatial frequency tuning across light levels (Fig 7d). We find that randomly pooling from these diverse classes of RGCs produces model V1 receptive fields with only mild shifts in spatial frequency tuning across light levels(Fig 7e), consistent with our experimental data (Fig 5c).

We explored how convergence impacts the shift in noise correlations at the periphery relative to V1. Each model V1 cell pools from units with covariance defined by their separation in retinal space (Greschner et al., 2011; Ruda et al., 2020), and difference in RGC cell class. There are several key features of the retinal data to consider when studying the convergence of these parallel pathways. First, RGC classes show diverse magnitudes and spatial extents of noise correlations across retinal space, as well as distinct degrees of modulation in noise correlation magnitude with light level. Second, RGCs of the same class, but opposite polarity, exhibit negative noise correlations (Greschner et al., 2011; Ruda et al., 2020). Finally, inter-polarity and inter-class correlations are much weaker and more retinotopically restricted than correlations between RGCs of the same class and polarity (Ala-Laurila et al., 2011; Greschner et al., 2011; Ruda et al., 2020). Given these prior results in the retina, we employed a simple model in which the activity of each RGC class was independent from all other classes (cross-class covariance=0).

We begin by modeling V1 neurons as pooling from a single class of RGCs, arranged in a mosaic, for which the change in retinal noise correlations across luminance conditions is clearest (Fig 7f, left). In the rodent retina, this corresponds to the OFF Brisk Transient RGCs, or ON Parasol RGCs in the primate (Greschner et al., 2011; Ruda et al., 2020). Given the regular spacing of RGCs in a mosaic and the diameter of V1 receptive fields, each model V1 neuron receives significant input from fewer than 10 afferents. In this model, neighboring V1 neurons have unrealistically large, positive noise correlations, and the difference in noise correlations across light levels is apparent (Fig. 7f, right). The cortical noise correlation distribution in this model represents a recapitulation of the retinal noise distribution when the responses of highly correlated, neighboring RGCs from a single class are pooled.

A more realistic model of the feedforward projections to V1 requires convergence of ON and OFF afferents (Hubel and Wiesel, 1962; Ferster et al. 1996). We next model the V1 population as pooling over an ON and OFF mosaic of a single class. We keep the total number of inputs to each V1 cell constant by pooling from each RGC within the receptive field with 50% probability. This random pooling across negatively correlated RGC populations decorrelates V1 activity relative to the prior model and results in more overlapping noise correlation distributions across light levels (Fig. 7g).

Adding additional RGC classes, with their specific magnitude and spatial extent of noise correlations, further reduces the noise correlations in model V1 neurons. Pooling across RGC classes leads to noise correlations that closely matching our data from mouse V1 (Fig 7h, i). The convergence of diverse retinal inputs onto a V1 neuron can therefore explain why the large differences in noise correlations across light levels observed in retina become undetectable in V1.

## Discussion

We have shown that visually evoked responses in mouse V1 are largely invariant to light adaptation state, despite changes in RGC and LGN receptive fields. Using both two-photon calcium imaging and electrophysiology, we found that spatiotemporal tuning of V1 neurons does not significantly shift with light level, as reflected in the consistent responses to both natural movies and gratings across light levels. In addition, we found that light level is not a strong determinant of noise correlation magnitude in mouse V1. In contrast, LGN responses, like their RGC input, shift from scotopic to photopic conditions: spatial frequency tuning shifts higher and noise correlation magnitudes decrease for nearby LGN cell pairs. We demonstrate that these discrepancies between retinal and cortical representations with luminance can be accounted for by the random convergence of functionally diverse peripheral afferents onto single cortical neurons.

The peripheral visual system adapts to the dramatic decrease in luminance in scotopic conditions by changing which photoreceptors are used, integrating over larger spatial extents and altering the relationship between neurons. These adaptations allow neurons to convey visual information, but alter the code by which that information is communicated. These changes in the peripheral representation pose a challenge to central processing areas, for which it is important that representations are invariant to light level. We demonstrate here that the representation in primary visual cortex is largely invariant to the changes at the periphery, and provide an explanation based on the integration of parallel functional pathways which converge in visual cortex.

This difference in spatial coding between peripheral and central sensory stations across luminance has been noted before. RGC receptive fields become more spatially lowpass at scotopic light levels due to growth of the receptive field center a reduction in the strength of the antagonistic surround (Barlow et al., 1957). In addition, RGCs integrate stimuli over a longer temporal window under scotopic conditions (Enroth-Cugell and Shapley, 1976). In contrast, our work and that of Duffy and Hubel (2007) have uncovered consistent selectivity in V1 across light levels.

While we have presented a simple, physiologically-constrained feedforward model to explain the invariant noise structure in V1, there may be additional, complimentary mechanisms that explain our experimental results. V1 excitatory cells receive a wide range of recurrent, feedback, and modulatory inputs in addition to that which is fed-forward from the retina. This may explain why cortical noise correlations *in-vivo* are more spatially extensive than predicted by our feedforward model, in which all noise correlations emerge due to overlapping feedforward receptive fields. These other sources of input may provide stronger sources of correlation than those present in the feedforward input. Indeed, for many V1 cells, the feedforward current from the LGN comprises less than 50% of the total current to a V1 cell (Finn et al., 2007; Li et al., 2013; Lien and Scanziani, 2013; Barbera et al., 2022). Thus, it is possible that any change in the feedforward component of the noise is masked by variations in other inputs that more strongly drive a V1 cell.

Cortical inhibition may also contribute by rapidly decorrelating excitatory cell activity such that the noise structure remains invariant despite changing noise correlations in the inputs. Studies of excitation-inhibition interactions in the cortical circuit have shown that inhibition, when coupled in time to excitatory fluctuations, can suppress noise correlations by subtracting off shared fluctuations in the circuit (Tetzlaf et al., 2012; Cardin, 2018). Inhibitory decorrelation of excitatory activity has been shown to enhance population encoding fidelity in olfactory bulb, somatosensory cortex, and retinal bipolar cells (Giridhar et al., 2011; Ly et al., 2012; Franke et al., 2017). In addition, pharmacologically blocking intracortical inhibition results in higher noise correlations between excitatory cells in somatosensory cortex (Sippy and Yuste, 2013). Parvalbumin-positive interneurons are particularly well suited to play a role in decorrelating excitatory cell activity by densely sampling activity within a local region of cortex-such that any shared fluctuations in excitatory activity, as would occur in a regime of highly correlated inputs, would drive strong inhibition form PV cells (Scholl et al., 2015). Fast inhibition mediated by these interneurons could dampen correlated activity fed-forward from the retina.

The results presented here are likely to generalize to primate V1. Across mammalian species, luminance evokes similar changes in retinal coding (Barlow et al., 1957; Enroth-Cugell and Shapley, 1973; Troy et al., 1999; Greschner et al., 2011; Grimes et al., 2014; Tikidji-Hamburyan et al., 2015; Ruda et al. 2020; Ruda et al., 2022). As we found in the mouse, the spatial selectivity of macaque V1 cells is largely invariant across scotopic and photopic conditions (Duffy and Hubel, 2007). Both species have parallel functional streams for visual processing that converge in visual cortex (O’Shea et al., 2023). It is therefore likely that the same mechanism of afferent convergence from LGN to cortex which explains our results in mouse V1 could account for invariant selectivity in the primate V1.

A fundamental challenge for sensory systems is to retain sensitivity across a broad range of environmental conditions without degrading selectivity for important features of the environment. The visual system must rapidly adapt to ambient luminance in order to maintain sensitivity. The shift between rod and cone mediated vision extends the retina’s dynamic range, but also alters RGC receptive fields and interneuronal correlations. These changes could disrupt downstream representations (Ruda et al., 2020; 2022). We propose that the key to maintaining a luminance-invariant representation in visual cortex is the convergence of signals across distinct afferent parallel pathways. Our proposal provides a mechanism by which the functional architecture of sensory systems, from peripheral receptors to cortex, can handle the dual challenges of maintaining sensitivity and selectivity across environmental conditions.

## Methods

### Animal Preparation

All experiments were approved by the University of Texas at Austin’s Institutional Animal Care and Use Committee, which maintains Association for Assessment and Accreditation of Laboratory Animal Care accreditation. All methods were carried out in accordance with relevant guidelines and regulations.

Imaging experiments were conducted using adult Ai94(TITL-GCaMP6s)-D;CaMK2a-tTA (Jackson Labs # 024115) mice of both sexes which express fluorescent calcium indicator GCaMP6s in forebrain excitatory neurons (n=10). Electrophysiology experiments were conducted using adult C57/BL6J (Jackson Labs #000664) mice of both sexes (n = 21).

For all surgical procedures, mice were anesthetized with isoflurane (2.5% induction, 1%– 1.5% surgery) and given two preoperative subcutaneous injections of analgesia (5 mg/kg carprofen, 3.5 mg/kg Ethiqa) and anti-inflammatory agent (dexamethasone, 1%). Mice were kept warm with a water-regulated pump pad. Each mouse underwent two surgical procedures.

The first was the mounting of a metal frame over the visual cortex using dental acrylic, used for head fixation of the mouse during experiments. The second was a 1-4mm diameter craniotomy drilled over V1 in the right hemisphere followed by a durotomy and a glass window implant. Surgical procedures were always performed on a separate day from functional imaging. Between the head frame implant and craniotomy, we mapped the V1 retinotopy with intrinsic signal imaging through the intact bone as described previously (Juavinett et al., 2017, Rhim et al., 2017). In all experiments we targeted the lower visual field of V1 based on the retinotopic map acquired with intrinsic signal imaging.

During all experiments, mice were awake, head restrained, and moved freely on a custom-built wheel. Mice were habituated to handling and head restraint over 1 week following recovery from the craniotomy in sessions increasing from 15 minutes to 1 hour.

### Imaging

Two-photon calcium imaging was performed with a Neurolabware microscope and Scanbox acquisition software. The scan rate varied between 10–15 frames/s, scaling with the number of rows in the field-of-view. A Chameleon Ultra laser was set to 920 nm to excite GCaMP6s. A Nikon x16 (0.8 NA 3 mm WD) or Olympus x10 (0.6 NA 3 mm WD) objective lens was used for all imaging sessions, with 900 μm and 1400 μm wide field of view respectively. All cells were imaged between 150 and 350 μm depth, which corresponds to layer 2/3 in the mouse. Average power of the exposed laser beam while imaging was approximately 60 mW and 30 mW for the x10 and x16 objective respectively.

Putative cells were extracted from our imaging data using the suite2p pipeline in Python (Pachitariu et al., 2017). We performed additional processing using custom MATLAB code to retain only cells meeting certain criteria. Cell fluorescent traces had to have skewness greater than 2. Cells also had to show significant visually evoked responses above baseline in their trial-averaged time courses as quantified by a one-sided, two-sample t-test with p<.001. Finally, only cells that were identified and retained for both scotopic and photopic conditions were analyzed. Cells were matched across conditions by finding ROIs matched in both location and morphology, using custom MATLAB code.

### Electrophysiology

The glass window over V1 was removed immediately prior to electrode implantation to expose the brain. All electrodes were implanted with a motorized drive (MP-285, Sutter Instruments). V1 recordings were made using a Neuronexus Buzsaki32 4 shank probe spanning 600 μm of cortex laterally, implanted acutely and lowered to 400-800 μm depth with electrode contacts spanning 140 μm vertically from the tip, or the Neuropixels 1.0 single shank probe, lowered to the same depth, with electrode contacts spanning the entirety of the probe. Raw traces were acquired at 30 kHz using the Ripple Grapevine Scout System for the Neuronexus probe, and the spikeGLX system for the neuropixels probe.

LGN recordings were made using a Neuropixels 1.0 probe implanted acutely and lowered to 2500-3200 μm, approximately 2500 μm posterior of bregma and 2000 μm lateral of midline. The electrode tip was lowered to the ventral extent of the LGN as identified by clearly evoked visual responses in the neural activity. Raw traces were acquired at 30 kHz using the spikeGLX System.

Putative single units were isolated from the raw traces offline using Kilosort3 (Pachitariu et al., 2023). We conducted further manual sorting of single units using the Phy GUI (https://github.com/cortex-lab/phy). All subsequent analysis of single unit spiking data was performed using custom MATLAB code.

### Visual Stimuli

#### Setup

A monochrome LED projector by Texas Instruments (Keynote Photonics) with spectral peak at 525 nm was used to generate stimuli with a 60Hz refresh rate onto a Teflon screen which provides a near-Lambertian surface (Rhim et al., 2021). The screen was 12.5 cm high x 32 cm wide, equating to approximately 64° x 116° of visual angle. Stimuli were coded using the Psychophysics Toolbox extension in MATLAB. Mice were positioned such that the perpendicular bisector from the mouse’s eye to the screen was 10cm with the screen angled at 30° from the mouse’s midline. The upper edge of the screen was placed near the vertical midline of the mouse’s field of view to target the lower visual field for stimulation.

#### Natural Movies

For both imaging and electrophysiology experiments, we presented two, 10 second long natural movies with each repeat followed by a 6 second long blank screen at mean luminance. Movie 1 consisted of honeybees flying in a garden (Ian Nauhaus, UT Austin) and Movie 2 consisted of monkeys playing in snow (David Leopold, NIMH).

#### Gratings

An identical set of full-field drifting sinewave gratings were used to characterize spatial and temporal tuning in LGN and V1. To measure spatial frequency tuning, gratings were presented at 5 spatial frequencies .0125, .025, .05, .1, .2 cycles/degree, all drifting at a temporal frequency of 2 cycles/sec, at 0 and 90 degrees, for 10 repeats of each stimulus. To measure temporal frequency tuning, gratings were presented at 6 temporal frequencies .5, 1, 2, 4, 8, 16 cycles/sec, all at a spatial frequency of .04 cycles/deg, at 0 and 90 degrees, for 10 repeats of each stimulus.

Static sinewave gratings used for V1 noise correlation experiments were presented at .05 cycles/° for 500ms at 4 orientations (0, 45, 90, 135°) and 2 contrasts (30%, 100%), each followed by a 1 second blank screen at mean luminance. Static sinewave gratings used for V1 orientation discrimination experiments were presented at .05 cycles/° for 500ms at 6 orientations (0, 10, 45,90, 100,135°) and 2 contrasts (30%, 100%), each followed by a 1 second blank screen at mean luminance. Drifting sinewave gratings used for LGN noise correlation experiments were presented at .01 cycles/° for 2 seconds at 4 orientations (0, 45, 90, 135°) and 2 contrasts (30%, 100%), each followed by a 1 second blank screen at mean luminance.

### Light Adaptation

At least 10 minutes prior to each experiment, the pupil was fully dilated with 1% atropine. Rod isomerization rates for each projector configuration were computed as previously described, using spectral radiance measurements taken with a PR655 spectroradiometer (Rhim et al., 2021; Rhim and Nauhaus, 2023). In all experiments, stimuli were presented first under scotopic conditions, then under photopic conditions.

#### Scotopic adaptation

Scotopic luminance was achieved by lowering projector power and adding two 1% neutral density filters (Thorlabs) to the light path. Care was taken to black out any other sources of visible light during the experiment, and supervision of the experiment was done remotely in an adjacent room separated by a shut door and floor to ceiling black out curtains. Based on our spectral radiance measurements, this configuration generated 25 Rh* rod^−1^ s^−1^. Mice viewed a blank screen at mean luminance for 30 minutes prior to any stimuli being played to achieve scotopic adaptation.

#### Photopic adaptation

We then raised the projector power and removed the neutral density filters to generate photopic luminance at 1.13 x 10^5^ Rh* rod^−1^ s^−1^. Mice viewed a blank screen at mean luminance for 10 minutes prior to any stimuli being played to achieve photopic adaptation.

### Analysis

#### Imaging

All cellular fluorescence traces were baseline normalized on a single-trial basis with the mean fluorescence value in the 200ms preceding stimulus onset. All visual responses were analyzed in units of percent change in fluorescence above baseline (Δ F/F). Δ F/F = [F(t)-p]/p, where F(t) is the raw fluorescence trace and p is the mean fluorescence magnitude in the baseline period. Signal correlations were computed as the zero-lag cross-correlation of the trial-averaged time courses for each cell, normalized by the autocorrelation of each cell.

For the orientation discrimination analysis, we compute each cell’s response on a single trial basis as the mean Δ F/F from 450 to 1300ms following stimulus onset, which captures the peak of the evoked fluorescence response to the 500ms static gratings for cells expressing GCaMP6s. The dDR procedure followed from Heller and David, 2022. For each unique pair of grating stimuli separated by 45°, we create 2 matrices containing the single trial responses for all simultaneously recorded cells. The difference between the trial-averaged means of these matrices gives the first axis in the dDR space-d╭. We then zero-center each of the response matrices and concatenate them, performing principal component analysis on this new matrix to find the first principal component of the zero-centered data (e_1_). The second axis in the dDR space is the component of e_1_ which is orthogonal to d╭ -the noise axis. The neural data matrices are then projected into the dDR space. In this 2-D space, we compute the discriminability index for a pair of stimuli (*d′*) as

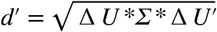

Where Δ *U* is a 2D vector of the difference in mean response to the two conditions, and Σ is the 2x2 covariance matrix.

To confirm the robustness of our chosen dimensionality reduction method, we compared *d* values computed in the dDR space to those computed for a low dimensional representation of our neural data generated with partial least squares (PLS) analysis (Rumyantsev et al., 2020). To directly compare our *d* values to previous reports of orientation discrimination in mouse V1 with 2-photon imaging, we followed the PLS procedure described by Rumyantsev et al. to generate a 5-dimensional representation of the neural data for each pair of stimuli, and computed *d* in the low dimensional space as above.

Noise correlations were computed for each pair of simultaneously recorded cells using the same procedure as Ruda et al., 2020. We first compute the cross-correlation function (CCF) on a single trial basis, normalized by the autocorrelation of each cell. Taking the mean of these single trial CCFs generates the “raw CCF.” Next, we shuffle all combinations of non-concurrent trials and compute CCFs for each of these combinations. Taking the average of this set of shuffled CCFs gives the “shuffled CCF.” We then subtract the shuffled CCF from the raw CCF to get the “noise CCF.” We take the value of the noise CCF at zero lag as our measure of noise correlation.

For plotting noise correlations as a function of lateral separation, we defined the location of each cell as the center of mass of the cell mask given by suite2p.

To model noise correlation as a function of lateral separation and r_sig_, we followed the model proposed by Smith and Kohn, 2008.

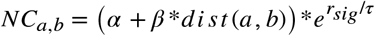

We grouped our noise correlation data into 100 bins corresponding to pairs with a given lateral separation and r_sig_. We took the mean of the noise correlation values in each bin and fit the model to these 100 data points using the fminsearch function in MATLAB to find parameter values that minimized the mean squared error of the fit. We computed the variance explained of the model fit as one minus the mean squared error divided by the variance of the data.

#### Electrophysiology

We computed the firing rate of each single unit in 50ms bins to get response time courses for all stimuli. We used these binned time courses to compute signal and noise correlations following the same procedure as described above for fluorescence time courses. For plotting noise correlations as a function of lateral or dorso-ventral separation, we defined the location of each single unit based on the electrode contact for which the spike waveform had the largest amplitude.

#### Tuning curves

Spatial and temporal frequency tuning curves for single units in LGN and V1 were fit to the trial-averaged evoked firing rate of each single unit. For each stimulus presentation trial, we sum all spikes, and divide by trial length (2 seconds) to get the firing rate in units of spikes/sec. By averaging the firing rate across all trials, we obtain the trial-averaged evoked firing rate for each stimulus. Baseline firing rate is computed in the same way, by taking the average firing rate during all times when the screen was blank. We presented all spatial and temporal frequencies at 2 directions (0 and 90°) and for each single unit determined the “preferred” direction as that evoking the maximum firing rate. Tuning curves were fit to responses at the preferred direction using a Gaussian function in the ln(spatial frequency) or ln(temporal frequency) domain using the fminsearch function in MATLAB to find parameter values that minimized the mean squared error of the fit. Before fitting, any firing rates below baseline were set to zero, and all firing rates were normalized to the maximum stimulus evoked response. Center-of-mass was computed as

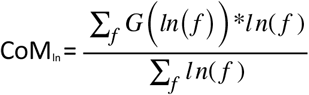

where G() is the Gaussian tuning curve and *f* is the stimulus domain in units of cycles/degree or cycles/sec. CoM is computed in the stimulus domain as CoM= *e*^*CoM*^*ln*.

### Visual pathway model

#### Modeling V1 receptive fields

We constructed a feed-forward model of V1 receptive fields following the statistical connectivity model as first proposed by Soodak for explaining how a population of orientation tuned V1 neurons can emerge from pooling of RGC receptive fields (Soodak 1987). The first layer consists of RGC mosaics with positions defining the center of circular, Gaussian receptive fields in retinotopic coordinates (). ON and OFF RGCs are positioned across retinal space following the pair-wise interaction point process model (PIPP), which results in appropriately separated receptive field centers (Eglen et al. 2005; Ringach 2007). Briefly, the model begins with randomly positioned ON and OFF receptive fields. Over many iterations, new candidate coordinate for each RGC are proposed at random, but only accepted if there is sufficient separation from all other receptive field centers. Over 100 iterations, this generates ON and OFF RGCs arranged in a mosaic pattern to evenly tile visual space. For each RGC type considered, we use PIPP to generate a mosaic independently of all other types.

We set the center radius of all RGC receptive fields of a given type to be equivalent, with radii across light levels following the results presented in Ruda et al. 2022. In this model, we do not consider the antagonistic surround of the RGC receptive field. Ruda et al. tracked receptive field size in 6 RGC populations across light levels.

| Type | Scotopic center radius ($\mu$ m) | Photopic center radius ($\mu$ m) |
| --- | --- | --- |
| ON brisk transient | 135 | 110 |
| OFF brisk transient | 140 | 130 |
| ON transient | 100 | 90 |
| OFF transient | 110 | 95 |
| ON brisk sustained | 150 | 145 |
| OFF brisk sustained | 110 | 105 |

As a simplified assumption, in this model, we assume that LGN provides a 1:1 relay from retina to cortex. Therefore, in the second layer, each V1 unit pools directly from the RGC mosaics. Each V1 unit is assigned a 2-dimensional cortical position (), which corresponds to a retinotopic position as defined by the retino-cortical magnification factor. This retinotopic position is the center of the V1 receptive field. Here, to model the mouse visual pathway, we approximate the magnification factor as 400╭m cortex / 1000╭m retina (Garrett et al., 2014). The receptive field of V1 unit *a* is defined as a linear combination of the *i* receptive fields in the first layer, with weights (*w*) defined by a circular Gaussian with radius (ϕ) corresponding to the average radius of mouse V1 receptive fields∼ 5 -7° (Ringach, 2007; Niell and Stryker, 2008).

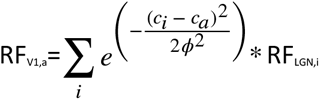

We compute the center-of-mass of these model V1 receptive fields from their Fourier spectrum.

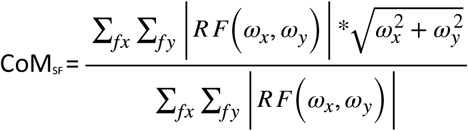

#### Modeling noise correlations

To model V1 noise correlations, we generate noisy RGC responses with each V1 unit pooling these responses with the synaptic weights defined as above.

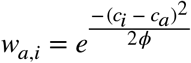

Where *w*_*a,i*_ is the weight of *i*th RGC input to V1 unit *a*.

We simulate noisy RGC responses to a repeated stimulus via random draws from a multivariate normal distribution using mvnrnd function in MATLAB. The mean response and variance of each model RGC can take an arbitrary positive value. For simplicity, we choose to assign equal mean and unit variance to all model RGCs (σ =1). The covariance is defined as a function of the distance between cell pair *i* and *j*:

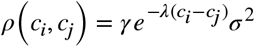

where γ scales the magnitude and λ scales the spatial extent of the covariance. In modeling the scotopic vs photopic states, only γ is varied, in keeping with the data presented in Ruda et al., 2020.

By simulating many instances of the RGC population response to generate simulated V1 responses, we estimate the noise correlations between model V1 cell pairs. In this modeling framework, we can also compute the noise correlation between V1 pair *a* and *b* as a function of their feedforward synaptic weights (*w*_*a*_ *w*_*b*_) and the covariance of their inputs .

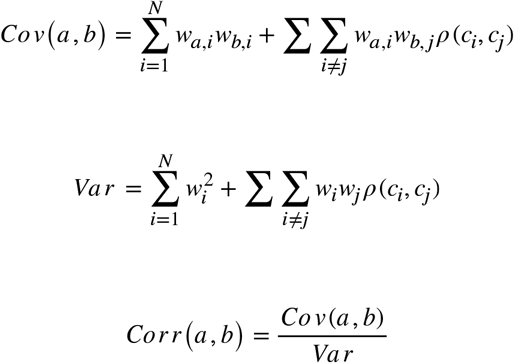

Each RGC type introduced into this model has a unique covariance matrix which is independent from all others. Below are the exponential function parameters used for the simulations of retinal covariance structure in Fig 7.

| $\rho$ | $\gamma$ scoto | λscoto | $\gamma$ photo | λphoto |
| --- | --- | --- | --- | --- |
| ON<sub>1</sub> – ON<sub>1</sub> | .5 | 6.5 | .15 | 6.5 |
| OFF<sub>1</sub> – OFF<sub>1</sub> | .5 | 6.5 | .15 | 6.5 |
| $ON_1 - OFF_1$ | -.1 | 8 | -.05 | 8 |
| $ON_2 - ON_2$ | .15 | 8 | .05 | 8 |
| $OFF_2 - OFF_2$ | .15 | 8 | .05 | 8 |
| $ON_2 - OFF_2$ | -.05 | 8 | -.025 | 8 |
| $ON_3 - ON_3$ | .25 | 6.5 | .2 | 6.5 |
| $OFF_3 - OFF_3$ | .25 | 6.5 | .2 | 6.5 |
| $ON_3 - OFF_3$ | -.075 | 8 | -.05 | 8 |

The vector of V1 weights *w* has the same retinotopic extent but is defined for additional RGC positions when more RGC types are added to the model. Changing the number of inputs to each V1 cell will have additional effects on the magnitude of V1 correlations. To avoid this confound when studying the effects of convergence in V1, we keep the number of inputs to each model V1 cell constant by drawing from a binomial distribution, distinct from the weights in *w*, such that each V1 cell pools randomly from a subset of RGCs within the retinotopic extent of its receptive field. The probability of pooling from any RGC declines with the addition of more RGC mosaics to the model.

| Model | Probability of pooling |
| --- | --- |
| Fig 7e | 1 |
| Fig 7f | 1/2 |
| Fig 7g | 1/4 |
| Fig 7h | 1/6 |

## Acknowledgements

The authors wish to thank Gabriela Coello-Reyes for essential guidance on surgical and experimental procedures, Carrie Barr for animal care and lab management, Charles Heller and Stephen David for instruction on the dDR method.

## Author contributions

All authors conceived and designed research. R.T.O. performed surgeries, performed experiments, and analyzed the data. R.T.O. prepared figures and drafted manuscript. All authors edited and revised manuscript.

## Data/code availability

Upon publication, data and code will be available on Figshare.

**Supplemental Fig. 1.**
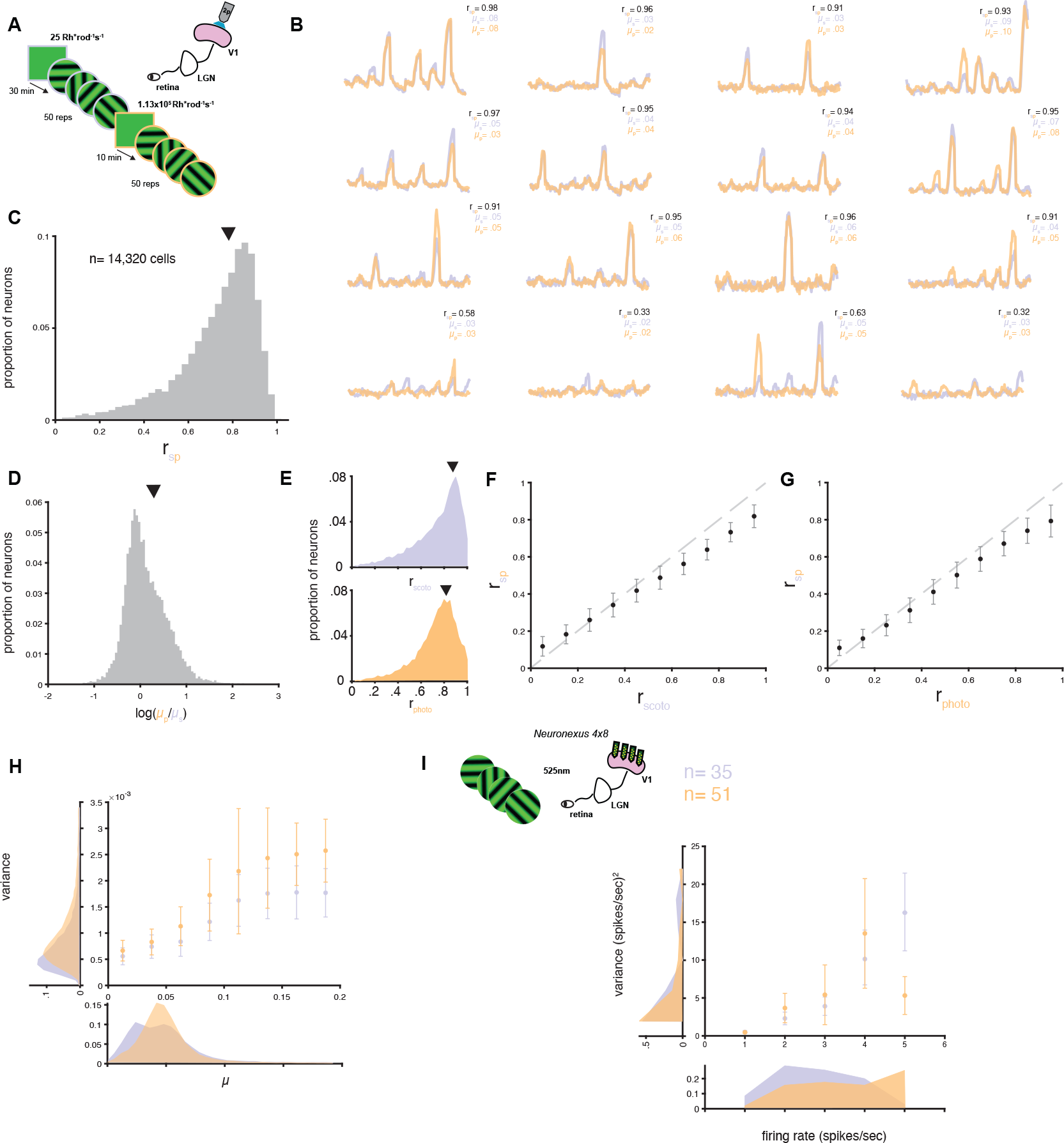
V1 responses to gratings in scotopic and photopic conditions: A. Experimental design. Awake mice were head fixed on a treadmill underneath a two-photon microscope while viewing 50 repeats of a 12-second series of static gratings at 4 orientations and 2 contrasts under scotopic (purple) and photopic (orange) conditions. To compare V1 responses under these two regimes, the same V1 neurons were tracked across conditions. B Trial-averaged, baseline normalized fluorescent responses (ΔF/F) of 16 example cells to the grating series at scotopic (purple) and photopic (orange) light levels. Inset gives the cross correlation of the mean responses across light conditions (r_sp_) and the mean over time of the baseline-normalized responses in scotopic (╭_s_) and photopic (╭_p_) conditions. The top 3 rows show example cell responses with high r_sp_. The bottom row shows example cell responses with low r_sp_. C. Histogram of r_sp_ for all cells tracked across luminance conditions. Arrow indicates median of distribution. D. Histogram of the log ratio of ╭_p_ /╭_s_ for all cells tracked across luminance conditions. Arrow indicates median of distribution. E. Histograms for within-condition correlation for trial-averaged responses taken from random splits of trials in scotopic (r_scoto_) (top) and photopic (r_photo_) (bottom) conditions. Arrows indicate median of distribution. F. r_sp_ plotted as function of r_scoto_, binned by within-condition correlation in 10 equally sized bins spanning 0 to 1. Scatter points indicate mean r_sp_ within each bin, bars indicate +/-.5 standard deviation. Gray dashed line indicates unity. G. r_sp_ plotted as function of r_photo_, binned by within-condition correlation in 10 equally sized bins spanning 0 to 1. Scatter points indicate mean r_sp_ within each bin, bars indicate +/-.5 standard deviation. Gray dashed line indicates unity. H. Response variance plotted as a function of response mean (╭) for all cells in scotopic (purple) and photopic (orange) conditions. Variance ((ΔF/F)^2^) binned by ╭ in 8 equally sized bins spanning 0 to .2 ΔF/F. Scatter points indicate average variance within each bin, bars indicate +/-.5 standard deviation. Marginal distributions of mean (bottom) and variance (left) for all cells in in scotopic (purple) and photopic (orange) conditions. I. (Left) Experimental design for electrophysiology recordings. Awake mice were head fixed on a treadmill with a 4x8 Neuronexus probe implanted in V1 while viewing 12-second series of static gratings at 4 orientations and 2 contrasts presented under scotopic and photopic light adaptation states. (Right) Response variance plotted as a function of mean firing rate (spikes/sec) for all cells in scotopic (purple) and photopic (orange) conditions. Variance ((spikes/sec)^2^) binned by mean in 5 equally sized bins spanning 1 to 5 spikes/sec. Scatter points indicate average variance within each bin, bars indicate +/-.5 standard deviation. Marginal distributions of mean (bottom) and variance (left) for all single units in in scotopic (purple) and photopic (orange) conditions.

**Supplemental Fig. 2.**
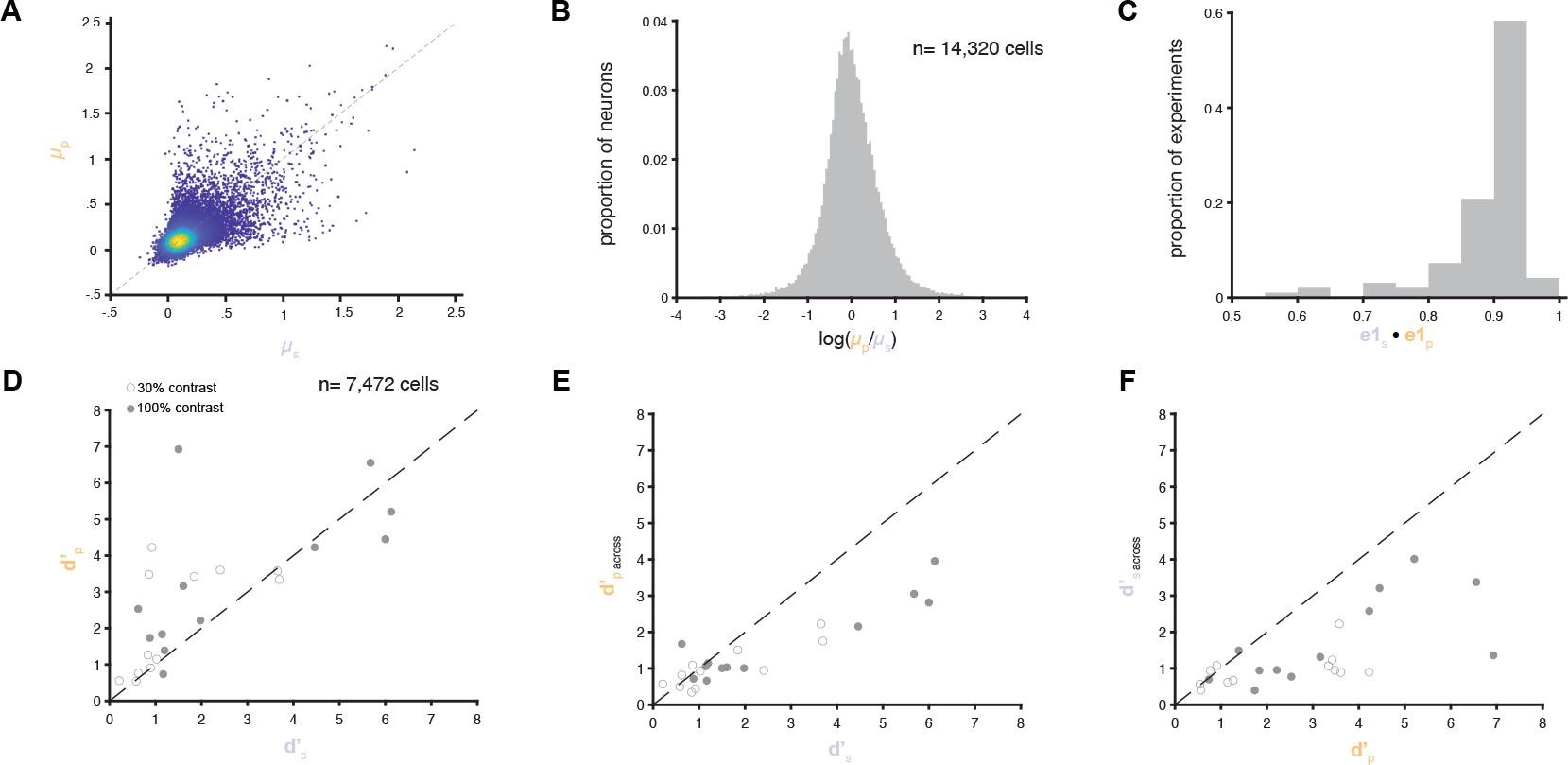
Discriminability of the V1 population code for orientation under scotopic and photopic conditions: A. Trial-averaged evoked responses (F/F) for all grating stimulus for each cell in scotopic (╭_s_) and photopic (╭_p_) conditions. Dashed line indicates unity. Color indicates density of scatter points (Robert Henson (2023). Flow Cytometry Data Reader and Visualization (https://www.mathworks.com/matlabcentral/fileexchange/8430-flow-cytometry-data-reader-and-visualization), MATLAB Central File Exchange). B. Histogram of the log ratio of ╭_p_ /╭_s_ for all stimuli and all cells tracked across luminance conditions. C. Histogram of the dot product of the first eigenvectors of the covariance matrix computed for stimulus pairs in the scotopic (e_1s_) and photopic (e_1p_) conditions, for the same V1 cells in each experiment. D. Scatter plot of the discriminability index computed for each pair of stimuli in each experiment independently for scotopic (d′_s_) and photopic (d′_p_) conditions using the PLS dimensionality reduction method. E. Scatter plot of the discriminability index computed for each pair of stimuli in each experiment using scotopic data to fit the PLS space into which both the scotopic and photopic data were projected to compute d′_s_ and d′_p, across_. F. Scatter plot of the discriminability index computed for each pair of stimuli in each experiment using photopic data to fit the PLS space into which both the scotopic and photopic data were projected to compute d′_p_ and d′_s, across_.

